# *Smad4* and TGF-β1 dependent gene expression biomarkers in conditional intestinal adenoma, organoids and colorectal cancer

**DOI:** 10.1101/2024.03.15.584288

**Authors:** Mirvat Surakhy, Julia Matheson, David Barnes, Emma J. Carter, Jennifer Hughes, Claudia Bühnemann, Sabina Sanegre, Hans Morreau, Paul Metz, Charlotte J. Imianowski, A. Bassim Hassan

**Affiliations:** Oxford Molecular Pathology Institute, Sir William Dunn School of Pathology, University of Oxford, South Parks Road, Oxford, OX1 3RE, United Kingdom; Department of Pathology, Leiden University Medical Centre, Leiden, The Netherlands; Institute of Cancer and Genomic Sciences, University of Birmingham, Birmingham B15 2TT

**Keywords:** Smad4, Apc, intestine, adenoma, TGF-β1, Id1, Spp1, Pak3, gene expression, colorectal cancer, biomarkers

## Abstract

TGF-β ligand activation suppresses cell growth yet can paradoxically and potently promote cancer invasion and metastasis depending on downstream pathway mutational context. Here, we evaluated the basis of this observation in conditional murine intestinal adenoma models with and without loss of Mothers against decapentaplegic homolog 4 (*Smad4*), with the aim of identifying TGF-β-BMP-SMAD4 pathway dependent gene expression biomarkers for translational application. Conditional *Lgr5*-CreER^T2^ activation in *Apc^fl/fl^Smad4^fl/fl^*resulted in adenoma formation with recombined homozygote floxed alleles *(Apc*^Δ/Δ^*Smad4*^Δ/Δ^). The adenoma phenotype was discordant, with a reduced small intestinal adenoma burden yet development of large non-metastatic caecal adenoma with nuclear localisation of phospho-Smad2/3. Derived *Apc*^Δ/Δ^*Smad4*^Δ/Δ^ adenoma organoids resisted TGF-β1 dose dependent growth arrest and cell death (IC_50_ 534pM) compared to *Apc*^Δ/Δ^*Smad4*^+/+^ (IC 24pM). TGF-β1 (390pM) modified adenoma mRNA expression (bulk RNA-Seq) most significantly for *Id1^low^* and *Spp1^high^* in *Apc*^Δ/Δ^*Smad4*^Δ/Δ^. Single cell RNAseq of caecal adenoma identified expansion of *Lgr5^low^, Pak3^high^* and *Id1^low^* progenitor populations in *Apc*^Δ/Δ^*Smad4*^Δ/Δ^. Of the 76 *Smad4* and TGF-β1 dependent genes identified in adenoma organoids, 7 human equivalent genes were also significantly differentially expressed in colorectal cancer, including *ID1^low^*, *SPP1^high^* and *PAK3^high^* that also correlated with poorer survival (TCGA cohorts). Murine conditional models identified *Smad4* loss of function mRNA expression biomarkers that require further evaluation as functional classifiers of colorectal cancer subtypes.

## Introduction

Mutation of the downstream TGF-β-BMP ligand-dependent signalling pathway can result in context-dependent and paradoxical gain of function (GOF) of the cancer phenotype, i.e. the ligand activation of the normally functioning signalling pathway exerts a tumour suppressive effect, yet following downstream pathway mutation, ligand activation can then result in ‘oncogenic’ GOF activation, epithelial-mesenchymal transition (EMT) and potent promotion of the invasive and metastatic cancer phenotype ^1^. It is well established that human cancer associated variants (mutations) of the TGF-β-BMP-SMAD4 pathway are correlated with poorer prognosis in enriched sub-types of intestinal cancers such colorectal and pancreatic adenocarcinoma ^2^. Despite the incorporation of whole genome sequencing, gene level RNA sequencing and proteomic biomarkers into supervised machine learning prognostic classifiers, for example in colorectal cancer, there remains a paucity of pathwayspecific gene expression biomarkers more closely linked to molecular mechanisms and indicative of pathway activity ^3,4^. This background provided the impetus for further evaluation of the critically important TGF-β-BMP pathway that when disrupted, is associated with the worse prognostic sub-types of intestinal cancer. Here, we use murine conditional *Apc* dependent intestinal models to generate *in vivo* and *in vitro* adenoma with and without TGF-β-BMP pathway modification to prospectively identify pathway and context specific gene expression.

The TGF-β1 ligand acts as a dimer to bring together specific heterodimeric cell surface type I and II serine/threonine kinase receptors to activate signals by phosphorylation of R-Smad transcription factors. For example, TGF-β1 binds to TGFBR1 and TGFBR2 to signal through p-Smad2/3, but can also bind to activin receptor type 1A (ACVR1) to transduce signal through p-Smad1/5 ^5,6^. Following receptor activation, phospho-R-Smads form either heterotrimeric or heterodimeric complexes with Smad4, the co-Smad, as it continuously shuttles between the cytoplasm and nucleus ^5,7^. Smad4-R-Smad complexes bind with cofactors and DNA to co-ordinate gene expression ^6^. Chromatin regions can alter quickly (within 10 minutes) following TGF-β1 stimulation, with assembled Smad complexes binding to *cis*-regulatory elements with AP-1 footprints ^8,9^. Smad4 and R-Smad proteins consist of two highly conserved globular domains, named Mad homology domains, both at the Nterminus (MH1) and C-terminus (MH2) separated by a less well-conserved serine and proline-rich linker region critical for protein-protein interactions and stability ^6,9,10^. The MH1 domain is involved in DNA binding through a conserved identical hairpin structure present in all members except Smad2 ^5,11,12^. The majority of SMAD4 loss of function (LOF) cancer somatic mutations are missense, although nonsense, splice site, frameshift, and inframe insertion/deletion mutations exist. In terms of protein localisation, frequent hot spot mutations occur within the conserved R-SMAD binding surface of the MH2 domain and missense mutations reported in the MH1 domain along the DNA-binding interface ^13^.

The specialised intestinal epithelium is comprised of crypt-villus functional cell units that provide an ideal experimental system for the investigation of specific components of the TGF-β signalling pathway ^14^. Differentiated cells in the crypt and villous comprise of four epithelial lineages: enterocytes, goblet, entero-endocrine and Paneth cells ^14^. Leucine-rich repeat containing G-protein-coupled receptor (*Lgr5*) positive crypt base columnar cells (CBC) are stem cells that interact with Paneth cells to maintain the stem cell niche through *Wnt* activation and β-catenin stability ^15,16^. *Lgr5* positive CBC cells divide, renew and create transit amplifying cells that undergo 4 to 5 cycles of cell division before differentiating as they ascend the villus. By using a *Lgr5* driven Cre recombinase, genes with associated loxP sites can be randomly disrupted in CBCs once the Cre is activated. BMP signalling is active at the crypt-villous border towards the villous tip and is controlled by a ligand gradient to promote differentiation towards secretory and enteroendocrine cells ^17–19^. BMP concentrations are regulated in the intestinal crypts by several different BMP antagonists, such as Gremlin, Noggin and Chordin-like1 ^20^. TGF-β type I/II receptors and ligands are present and expressed in the muscularis, lamina propria and the differentiated compartment of both the small intestine and colon ^21^. TGF-β is also frequently present in the intestinal tumour microenvironment and is released either by cancer cells, tumour stromal cell or immune cells ^22^. Intestinal adenoma can be conditionally generated when *Apc* loss of function no longer regulates the degradation of β-catenin, resulting in constitutive activation of transcription by β-catenin/ Tcf4. Subsequent mutational events are selected for during the progression of adenoma to invasive carcinoma in humans, e.g. *RAS*/*MAPK* GOF, *PTEN* and *TP53* LOF, and later acquired LOF mutation of the TGF-β-BMP ligand activated receptor pathway, such a mutation of receptors and Smad2/3/4 ^23^. The conditional *Apc*^fl*/*fl^ model using *Lgr5CreER^T2^* is well characterised and so we incorporated loxP dependent (LOF) of *Smad4* (*Smad4*^fl*/*fl^ alleles) into this model.

Here, we first set out to evaluate loss of the TGF-β pathway signal transduction regulator, Mothers against decapentaplegic homolog 4 (*Smad4*), *in vivo* and *in vitro*, with respect to a murine intestinal adenoma phenotype and dependent gene expression. In view of the complexity of the interacting pathways that can alter TGF-β1-*Smad4* induced phenotypic effects, especially in immortalised human cancer cell lines, we sought to better define the contextual impact of *Smad4* LOF to specifically evaluate the *Smad4* and TGF-β1 ligand dependency on gene expression biomarkers and to translate these findings with respect to human intestinal cancer.

## Results

### Combined Apc and Smad4 conditional disruption results in discordant adenoma phenotypes

To investigate the genetic interactions between *Apc* and *Smad4* in the murine intestine, we combined conditional alleles with a *Cre* driven by the promoter *Lgr5* and a ROSA26-loxSTOP reporter allele (*Lgr5-EGFP-IRES-CreER*^T2^ or *Lgr5-CreER*^T2^)^24^ (Fig. 1a). Heterozygote alleles were inter-crossed (*Apc*^fl/+^*Smad4*^fl/+^*Lgr5-CreER^T2^*) on a C57Bl/6J background (backcrossed to C57Bl/6J for >10 generations) to generate homozygotes and heterozygote littermates that differ in Smad4 conditional alleles ^25,26^. Conditional recombination followed tamoxifen injection (4OHT) in adult animals (Fig. 1a). The floxed allele of *Smad4* (*Smad4*^Δ/Δ^) resulted in loss of the MH1 DNA binding domain and the production of a truncated 43kDa protein which was not capable of translocating to the nucleus (see below, Supplementary Fig. 1). The germline homozygote loxP alleles are denoted *Smad4*^fl/fl^ and *Apc*^fl/fl^, and where somatic tissues have been directly tested for the presence of the resulting Cre-induced recombined alleles (floxed) using PCR, were denoted as *S m a d*^Δ/^*4*^Δ^ and *Apc*^Δ/Δ^. Recombination rates in intestinal epithelial cells were lower overall with the *Lgr5* transgene (5-6%), but highly specific for *Lgr5* CBC stem cells ^15,27^. Rapid adenoma development occurred as expected (median 35-40 days post injection 4OHT) and animals were culled if they exceeded predefined humane endpoints. Comparison of overall survival (Kaplan-Meier) between *Apc*^fl/fl^*Smad4*^fl/fl^*Lgr5-CreER* ^T2^ and *Apc*^fl/fl^*Smad4*^fl/+^*Lgr5-CreER*^T2^ with *Apc*^fl/fl^*Smad4*^+/+^*Lgr5-CreER*^T2^ controls showed a modest but significant improvement in survival (p=0.011, Fig. 1a). These observations suggest that introducing *Smad4* LOF in *Apc* LOF intestinal adenoma does not result in overwhelming tumour progression and metastasis. However, analysis of intestinal adenoma distribution between the small intestine and colon showed a marked discordance in growth phenotypes in *Apc*^fl/fl^*Smad4*^fl/fl^*Lgr5-CreER*^T2^. A significant reduction in small intestinal adenoma burden was observed with paradoxical growth promotion of adenoma in the caecum (Fig. 1b). As the distribution of adenoma on a C57BL/6 background is predominantly in the small intestine rather than the colon and caecum, the multiple small intestinal adenoma would be expected to promote anaemia, small intestinal obstruction and decreased survival ^27,28^. The significantly reduced small intestinal adenoma burden in *Apc*^fl/fl^*Smad4*^fl/fl^*Lgr5-CreER*^T2^ compared to controls may have improved overall survival if this was the only phenotypic effect (Fig. 1a, b). Examination of the caecum and large intestine revealed markedly enlarged coalesced adenoma where numerical analysis of single adenoma was impossible, hence we reported caecal weight as a surrogate (Fig 1b). The adenoma burden in heterozygote *Apc*^fl/fl^*Smad4*^+/fl^*Lgr5-CreER*^T2^ compared to controls was statistically non-significant and does not support either *Smad4* haplo-insufficiency or a dominant negative effect of *Smad4*^fl/+^. Histological grading of small intestinal adenoma showed low-grade dysplasia in all genotypes, whereas more aggressive carcinoma *in situ* was observed in regions of *Apc*^fl/fl^*Smad4*^fl/fl^*Lgr5-CreER*^T2^ in the caecum (Fig. 1c, author HM pathological review). Examination of H&E stained small intestinal adenoma tissue sections showed that *Apc*^fl/fl^*Smad4*^+/+^ mice had sessile adenomatosis appearance compared to *Apc*^fl/fl^*Smad4*^fl/fl^ that had mainly tubular adenomas. Crypt abscesses were detected in both genotypes but no tissue invasion and metastasis were observed (Fig. 1c). The efficiency of Cre-mediated *Smad4* allele deletion was examined by immunohistochemistry (IHC, Fig. 1c). Smad4 protein was clearly detected in *Apc*^Δ/Δ^*Smad4*^+/+^ adenoma and adjacent tissue as expected. Adenoma from *Apc*^Δ/Δ^ *Smad4*^Δ/Δ^ did not label strongly with anti-Smad4 antibodies, consistent with adenoma-specific disruption in both the small and large intestine (Fig.1c). The presence of active TGF-β (BMP) signaling was supported by p-Smad2 and p-Smad1/5/8 nuclear immunolabeling, respectively, as was confirmed by immunoblots (Fig. 1d). Both p-Smad2 and p-Smad 1/5/8 labelling were detected in adenoma from *Apc*^Δ/Δ^*Smad4*^Δ/Δ^ compared to normal tissue and littermate control adenoma, consistent with either TGF-β alone or with BMP ligand activation in combination (p<0.001, Fig. 1d).

**Figure 1.**
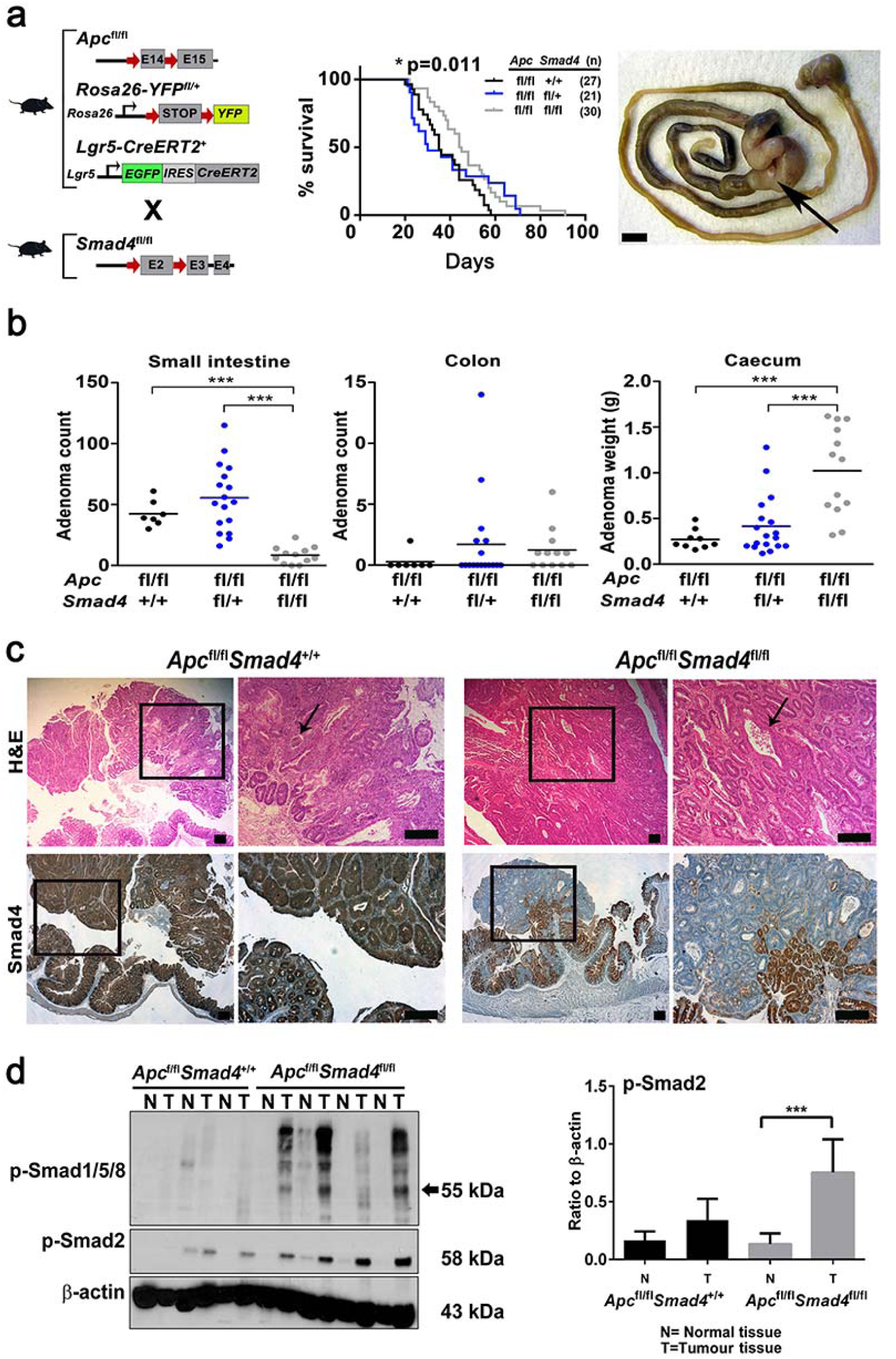
Adenoma growth following conditional disruption of Apc and Smad4 loxP alleles using Lgr5-CreER a. Breeding strategy to combine loxP conditional alleles with *Lgr5-CreER*^T2^ prior to 4OHT injection in adult mice. Kaplan-Meier survival of *Apc*^fl/fl^*Smad4*^+/+^, *Apc*^fl/fl^*Smad4*^fl/+^ and *Apc*^fl/fl^*Smad4*^fl/fl^ mice post-Cre induction. Comparison between *Apc*^fl/fl^*Smad4*^+/+^ and *Apc*^fl/fl^*Smad4*^fl/fl^ genotypes, *p=0.011, Log-rank test. Caecal enlargement (arrow) from *Apc*^fl/fl^*Smad4*^fl/fl^ mice. Bar 1cm. b. Caecum (adenoma) weight, small intestine and colon total adenoma from *Apc*^fl/fl^*Smad4*^+/+^ (n=9), *Apc*^fl/fl^*Smad4*^fl/+^ (n=17) and *Apc*^fl/fl^*Smad4*^fl/fl^ (n=13). Horizontal bars; mean. One-way ANOVA with Bonferroni’s multiple comparison test. ***p<0.001. c. Haematoxylin and Eosin (H&E) sections and anti-Smad4 antibody immunohistochemistry localisation in *Apc*^fl/fl^*Smad4*^+/+^ and *Apc*^fl/fl^*Smad4*^fl/fl^ caecal adenoma. Arrows show crypt abscesses, upper panel. Smad4 localisation is reduced in adenoma and confined to adjacent normal colonic crypt cells in *Apc*^fl/fl^*Smad4*^fl/fl^ mice. Scale bar 100µm. d. Western blot of adenoma (T) and normal tissue (N) from *Apc*^fl/fl^*Smad4*^+/+^ and *Apc*^fl/fl^*Smad4*^f/fl^ with anti-phospho-Smad2 (p-Smad2) and anti-phospho-Smad1/5/8 antibodies. Densitometric analysis for p-Smad2 represented as ratio to β-actin (for *Apc*^fl/fl^*Smad4*^+/+^ n=3 and for *Apc*^fl/fl^*Smad4*^fl/fl^ n=4 experimental replicates). Significant increase in p-Smad2 in adenoma (T) compared to normal tissues (N) in *Apc*^fl/fl^*Smad4*^fl/fl^ mice. ***p<0.001. Non-parametric one-way ANOVA with Dunn’s multiple comparison test. Mean ± S.D.

### Combined Apc and Smad4 disruption modifies growth of adenoma organoids

In order to generate an organoid culture, we isolated *Apc*^Δ/Δ^*Smad4*^Δ/Δ^ (*Smad4*^Δ/Δ^) and *Apc*^Δ/Δ^*Smad4*^+/+^ (*Smad4*^+/+^) caecal adenoma and generated *in vitro* cultures in Matrigel using published conditions ^24,29^. Using a ROSA26-LSL-YFP reporter line bred with *Apc*^fl/fl^*Smad4*^fl/fl^*Lgr5-CreER*^T2^, we confirmed that 40HT exposure resulted in uniform YFP expression in adenoma organoid culture that was also TGF-β1 independent (Fig. 2a). In culture, *Smad4*^Δ/Δ^ adenomas appeared smaller and grew slower compared to *Smad4*^+/+^ adenomas, as measured by both area and the alamarBlue (AB) growth assay (Supplementary Fig. 2). Noggin is a BMP pathway inhibitor that normally prevents enterocyte differentiation and is required for crypt culture maintenance ^23,24,29^. Noggin addition had no effect on adenoma growth (Supplementary Fig. 2). SB-505124, a TGF-βR1 small molecule inhibitor also had no significant inhibitory effect on Smad4^+/+^ and *Smad4*^Δ/Δ^ adenoma growth (1 μM) without the exogenous addition of TGF-β1 (Supplementary Fig.2). These data suggest that the adenoma culture conditions appeared to be independent of endogenous TGF-β supply. To exclude whether there may have been a genotype dependent difference in the floxing efficiency between *Apc* and *Smad4* that may have accounted for the two-fold differences in growth in culture, we also generated adenoma with an alternative conditional Cre (*Villin-CreER^T2^*). Here, cultured small intestinal organoids from healthy animals bred with non-floxed alleles, *Apc*^fl/fl^*Smad4*^+/+^*Villin-CreER^T2^Rosa26-YFP* and *Apc*^fl/fl^*Smad4*^fl/fl^*VillinCreER^T2^Rosa26-YFP* genotypes, were cultured in organoid media containing R-spondin-1, Noggin and EGF. Intestinal organoids that formed a normal crypt-villous morphology were exposed to 4OHT followed by withdrawal of R-spondin-1. This resulted in adenoma organoid formation in both *Apc*^Δ/Δ^*Smad4*^+/+^ and *Apc*^Δ/Δ^*Smad4*^Δ/Δ^ (Supplementary Fig. 3). Allele recombination was confirmed by PCR product of 259bp and 512bp for *Apc* and *Smad4 alleles*, respectively. Both the *Apc*^fl/fl^ and *Smad4*^fl/fl^ alleles recombined at the same time and rate following 4OHT treatment, and were detected as early as day 2 post treatment (Supplementary Fig. 3). Thus, floxing efficiency appeared similar between alleles.

**Figure 2.**
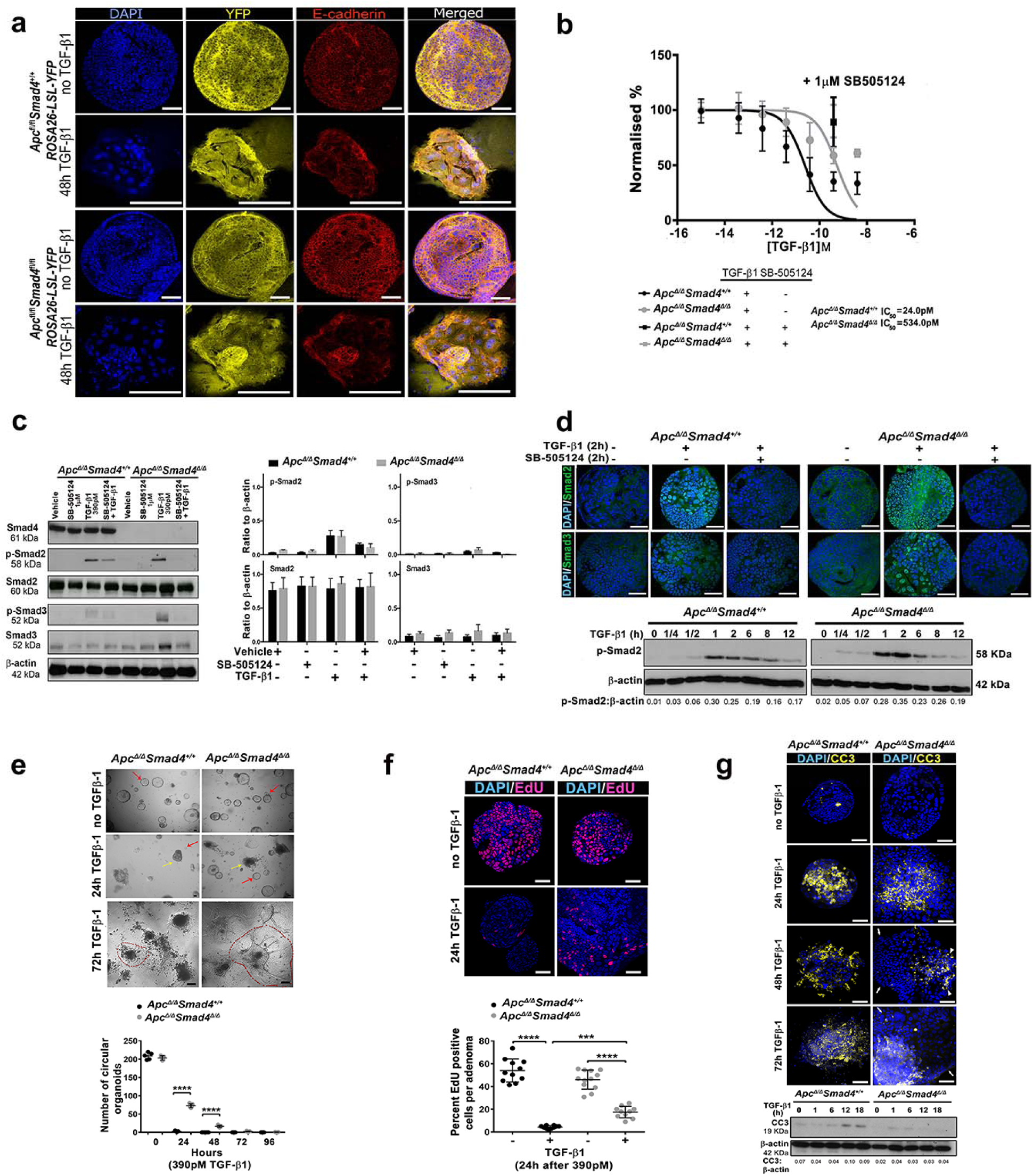
TGF-β1 and Smad4 dependent signalling in intestinal adenoma organoids. a. Expression of YPF reporter, E-cadherin localisation and DNA (DAPI) in *Apc*^Δ/Δ^*Smad4*^+/+^ and *Apc*^Δ/Δ^*Smad4*^Δ/Δ^ adenoma organoids with and without exposure to TGF-β1 (390pM). b. TGF-β1 (range: 0.0393900pM) induced growth inhibition of adenoma organoids quantified by alamarBlue (measured at 24h. Data normalised to untreated control (100%) (n=4 experimental replicates). Growth inhibition of 390pM TGF-β1 rescued by pre-treatment with SB-505124 (ALK4,5,7 inhibitor) for 2h (1mM). Error bars ± S.D. c. Representative immunoblots of Smad4, total RSmads and p-R-Smads from *Apc*^Δ/Δ^*Smad4*^+/+^ and *Apc*^Δ/Δ^*Smad4*^Δ/Δ^ adenoma organoids with and without SB-505124. Graphs of p-Smad2/ Smad2 and p-Smad3/ Smad3 densitometry relative to β-actin (n=3 experimental replicates). Nonparametric one way-ANOVA with Dunn’s multiple comparison. d. Confocal immunofluorescence images of Smad2 and Smad3 nuclear localisation (DAPI) following TGF-β1 (390pM) exposure and SB-505124 inhibition. Scale bar 50µm. Associated Immunoblots of p-Smad2 at different time points (h) following TGF-β1 (390pM), densitometry relative to β-actin. e. Bright field confocal images of organoids 24-72 hr following TGF-β1 exposure. Red arrows intact spherical (circular) organoid morphology, yellow arrows show shape change, red dotted line indicates organoids with marked disruption, cell spreading and cellular transformation. Quantification shows relative preservation of spheroid (circular) organoids in *Apc*^Δ/Δ^*Smad4*^Δ/Δ^. One-way ANOVA with Sidak’s multiple comparison test ****p<0.0001. f. Suppression of organoid proliferation following TGF-β1 treatment assessed with EdU incubation for 2h. Confocal images of EdU incorporation (green) in nuclei (DAPIblue) from *Apc*^Δ/Δ^*Smad4*^+/+^ and *Apc*^Δ/Δ^*Smad4*^Δ/Δ^ and quantification using ImageJ (n=10 organoids). One-way ANOVA with Sidak’s multiple comparison test *** p<0.001, ****, p<0.0001, respectively. g. Cleaved Caspase 3 (CC3) activation following 24-72 hrs of TGF-β1 (390pM). CC3 (arrowheads) labelling in confocal images supported by immunoblots. Arrows indicate margin of organoids. Scale bar 50µm.

### Deletion of MH1 domain of Smad4 following Cre-mediated recombination

To evaluate the expression of *Smad4* mRNA in *Smad4*^Δ/Δ^ and *Smad4^+/+^ in vitro*, we used bulk RNA-seq of adenoma organoids and discovered two different *Smad4* mRNA transcripts*. Smad4* gene tracking with Cuffdiff revealed transcripts from the intact *Smad4* gene, and a second transcript from the *Smad4* gene with exon 2 excised following loxP mediated recombination (Supplementary Fig. 1). Immunoblots confirmed expression of Smad4 full-length protein (61kDa) in both *Smad4^+/+^* and *Smad4*^+/Δ^ adenomas, with a smaller band of (43kDa) detected in protein extracts from *Smad4*^+/Δ^ and *Smad4*^Δ/Δ^ adenomas (Supplementary Fig. 1). *In silico* analysis revealed an open reading frame in exon 5 of the *Smad4* sequence and mapping with ExPASy revealed that exon 2 deletion results in a truncated protein in which the *Smad4* MH1 DNA binding domain is selectively deleted. The predicted size of the truncated protein was 43kDa, comprising the linker and Smad4 MH2 domain (Supplementary Fig. 1). The full-length Smad4 protein in *Smad4^+/+^* adenoma showed nuclear localisation following exposure of 390pM TGF-β1 for 2 hours. Under the same conditions, no nuclear localisation of Smad4 (43kDa) was detected in *Smad4*^Δ/Δ^ adenomas, consistent with loss of DNA binding function (Supplementary Fig. 1). Taken together, Cre-mediated recombination of the Smad4 gene resulted in selective MH1 deletion and a truncated protein that did not undergo nuclear localisation following TGF-β1 stimulation. These data, and the apparent reduction of detection using the same anti-Smad4 antibody in IHC of *Smad4*^Δ/Δ^ adenoma, are consistent with non-DNA binding and presumably soluble truncated SMAD4 protein that depletes from of cells during tissue processing resulting in reduced detection.

### TGF-*β*1 dose dependency in Smad4 ^Δ/Δ^ adenoma organoids

We next tested the dose and timing effects of exogenous addition of TGF-β1 to adenoma organoid cultures and the subsequent localisation of R-Smads. TGF-β1 induced dose-dependent growth inhibition of both the *Smad4*^+/+^ and *Smad4*^Δ/Δ^ adenomas, but with a 20-fold differential sensitivity (Fig. 2b). *Smad4*^+/+^ adenomas were more sensitive to TGFβ1, with an IC_50_ of 24pM and threshold of 0.39pM. For *Smad4*^Δ/Δ^, the TGF-β1 IC_50_ was 534pM with a threshold between 39-390pM. The decrease in adenoma viability with TGFβ1 dosing was prevented when the adenoma cultures were pre-incubated with SB-505124 (Fig. 2b). For subsequent experiments, a TGF-β1 concentration of 390pM was used to differentiate the TGF-β1 context-dependent sensitivity effects between *Smad4*^+/+^ (100 fold higher than the threshold) and *Smad4*^Δ/Δ^ (at threshold but below IC_50_) adenomas. TGF-β1 addition (390pM) resulted in the detection of phosphorylated p-Smad2 and p-Smad3 in both the *Smad4*^+/+^ and *Smad4*^Δ/Δ^ adenomas as expected (Fig. 2c). Despite the differential sensitivity, the kinetics of response to TGF-β1 stimulation were similar between genotypes, with p-Smad2 maximally detected after 1-2 hours and associated with nuclear localisation (Fig. 2c). Within 24 hours of TGF-β1 administration, >90% of the *Smad4*^+/+^ adenoma cells either had either morphological changes of apoptosis with organoid retraction, or changes in shape toward a spindle or mesenchymal phenotype (Supplementary Movies 1-4). These features appeared delayed to 72 hours in *Smad4*^Δ/Δ^ adenomas but were still detectable (Fig. 2e). The reduced viability and altered morphology were accompanied by a significant decreased in proliferation and increased apoptosis in *Smad4*^+/+^ adenoma when quantified by EdU and cleaved caspase 3 (CC3) labelling, respectively (Fig. 2f, g).

### TGF-*β*1 and Smad4-dependent Id1 and Spp1 gene expression in adenoma organoids

To assess the transcriptome response to TGF-β1 that might inform differences between *Smad4*^+/+^ and *Smad4*^Δ/Δ^ adenoma organoids, bulk paired-end RNA-seq detected 16,143 gene transcripts that we utilised for the analysis of differentially expressed genes (DEG). Three separate adenoma biological replicates per genotype and time were harvested at t=0, t=1 and t=12 hours after exposure to a 390pM TGF-β1 (Fig. 3a). Principle component analysis (PCA) of gene expression showed a clear separation of the samples based on genotype and duration of TGF-β1 exposure (Fig. 3b). Comparison of DEG between *Smad4*^Δ/Δ^ and *Smad4*^+/+^ at t=0 showed 111 upregulated genes (log_2_FC >1.5) and 51 downregulated genes (log_2_FC < −1.5) (Fig. 3c, Supplementary Table 1). The most important observation was the magnitude of the genotype dependent expression of a group of genes that included *Id1* (Inhibitor of DNA binding 1)*, Ddit4, Serpine1, Skil, Mucins, Myc, JunB* and *E2f* target genes (Fig. 3c, d, Supplementary Fig. 4). Gene ontology (GO) analysis of biological processes showed enrichment for epithelial cell proliferation and response to TGF-β1 (Supplementary Fig. 4, Supplementary Table 2). Gene set enrichment analysis (GSEA) using hallmark gene sets with “tmod” in *Smad4*^Δ/Δ^ showed downregulation of *Myc* target genes, differential regulation of TGF-β signalling genes, mainly by upregulation, and enrichment of other pathways including IL2-Stat5 and TNF signalling (AUC >0.5, padj<0.0001, Fig. 3e, Supplementary Table 2). Importantly, mucins in *Smad4*^Δ/Δ^ adenomas were among the DEGs with *Muc4, Muc20, Muc2* being upregulated and *Muc6* being downregulated (padj p<0.05, Supplementary Table 1, supplementary Fig. 5). Subsequent Mucin 2 immunolabeling confirmed protein overexpression in *Smad4*^Δ/Δ^ adenoma tissue sections (Supplementary Fig. 5).

**Figure 3.**
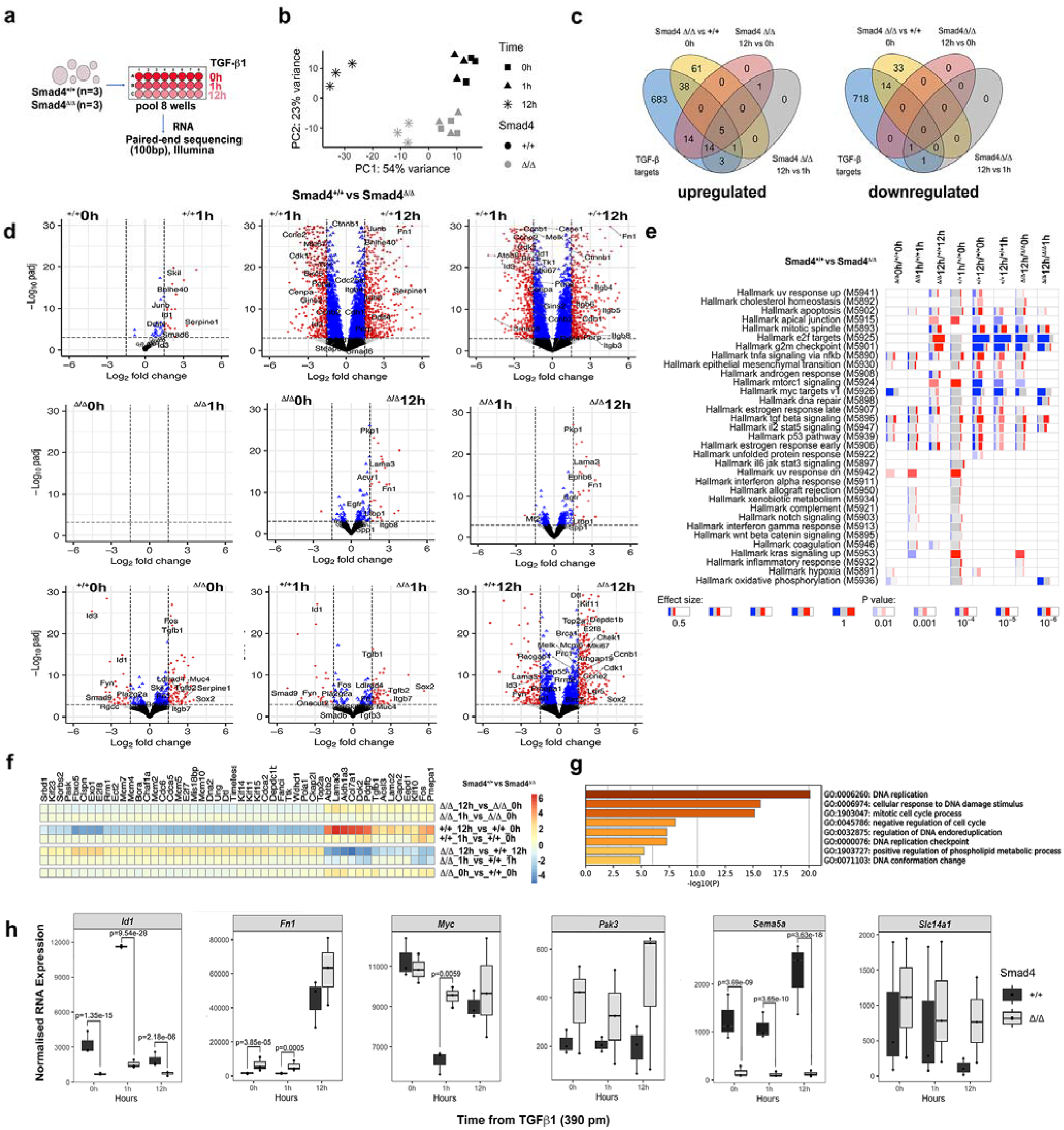
TGF-β1 and Smad4 dependent bulk RNA expression signatures in adenoma organoids. a. Adenoma organoids derived from 3 mice per genotype of *Apc*^Δ/Δ^*Smad4*^+/+^ and *Apc*^Δ/Δ^*Smad4*^Δ/Δ^ were exposed to TGF-β1 (390pM) (0, 1 and 12h, eight wells per time point), were then pooled for paired-end bulk RNA-Seq (total of 18 RNA-Seq samples). b. Principal components of transcriptional clusters according to genotype and time (vst-transformed). c. Venn diagram classifies TGF-β1 and Smad4 dependent down-regulated and upregulated genes. Threshold utilised 1.5 log2FC and adjusted p-value <0.05. d. Volcano plots of differentially expressed genes comparing time and genotype (^+/+^=*Apc*^Δ/Δ^*Smad4*^+/+^, ^Δ/Δ^ *= Apc*^Δ/Δ^*Smad4*^Δ/Δ^). In all volcano plots, genes with -log(p-adj)=30 are shown. Cut-off values for log2FC (1.5) and adjusted-pvalue (1e-3) are marked with dashed lines. *Apc*^Δ/Δ^*Smad4*^Δ/Δ^ DEGs are much reduced in number following TGF-β1 treatment. e. Enriched gene modules of DEG using the hallmark gene set enrichment (GSEA with tmod package). Comparisons shown in each column. Red and blue indicate the proportion of genes in a module that are either upregulated or downregulated, respectively. Width of each box relates to its effect size, where lighter, less saturated colours indicate lower significance with pvalues. f. Heatmap showing time-dependent changes in expression (log2FC) for the 50 genes with the lowest adjusted p-values (Likelihood ration test) and g. pathway enrichment for these genes are shown generated using Metascape. h. Summary of gene specific expression comparing *Apc*^Δ/Δ^*Smad4*^+/+^ and *Apc*^Δ/Δ^*Smad4*^Δ/Δ^ with time post TGF-β1 exposure. Statistical comparison performed using the Wald test in Deseq2, followed by Benjamini and Hochberg multiple hypothesis correction.

For early timepoints, DEG were mainly observed in *Smad4*^+/+^ adenoma organoids and included *Skil*, *Junb*, *Serpine 1*, *Ddit4* and *Id1* (Fig. 3d, Supplementary Fig. 4, Supplementary Table 1). The differential suppression of *Id1* in *Smad4*^Δ/Δ^ compared to *Smad4*^+/+^ after 1 hour achieved a 2.83 log_2_FC difference. By 12 hours, *Id1* maintained suppression (1.2 log2FC, Fig. 3d, Supplementary Table 1). Despite the repression of expression, Id1 protein abundance and immunolocalisation did not appear to be significantly altered between adenoma genotypes (Supplementary Fig. 6). We also observed that *Spp1* (Secreted phosphoprotein 1), an extracellular matrix gene, was the only gene significantly upregulated gene following TGF-β1 exposure (12 hours) in *Smad4*^Δ/Δ^ (log FC >1.5 threshold, Fig. 3c, d., Supplementary Table 3).

Later time points displayed further significant time dependent differences in gene expression. For example, 35 DEGs were observed in *Smad4^Δ/Δ^*compared to 689 in *Smad4*^+/+^ adenomas at t=12 hours vs t=0 hours (Fig.3d, Supplementary Table 1). Pathway enrichment analysis using tmod also showed that *E2f*, *G2m*, *Myc* and mTORC targets were more significantly downregulated in *Smad4*^+/+^ adenomas compared to *Smad4*^Δ/Δ^ adenomas (Fig. 3e, Supplementary Table 2). Time series analysis using the likelihood ratios showed differences in expression of genes including *Fos*, *Lama3, Aldh1a3, Col7a1* and *Doks* associated with increased induction by t=12 hours, whereas *E2f8, Exo1, Clspn, Kif14* and *Kif12* all were decreased in *Smad4*^+/+^ adenomas. Overall, the suppression of cell cycle genes was evident in *Smad4*^+/+^ compared to *Smad4*^Δ/Δ^ adenomas, as shown by Metascape overrepresentation (Fig.3f, g, Supplementary Table 3). Finally, TRRUST (Transcriptional Regulatory Relationship Unravelled by Sentence based Text mining) enrichment analysis implemented in Metascape also suggested potential regulation by transcription factors, with *Smad4*dependent downregulated genes being co-regulated by Sp1, Smad3 and Jun, and downregulated genes by Rbl2, Tp53 and E2f1 (Supplementary Fig.4). Several gene signatures were therefore determined that were *Smad4* and TGF-β1 dependent, with the two genes with the greatest magnitude of differential expression being *Id1* (low) and *Spp1* (high).

### TGF-*β*1 dependent gene expression and protein correlations

*Smad4* LOF did not fundamentally alter the range of TGF-β1 phenotypic responses, but did change the overall magnitude and relative gene expression. Whilst TGF-β1 administration in *Smad4*^+/+^ adenomas resulted in a >4.5 Log FC relative increase in expression *of Fn1*, we did not observe a similar change in Fn1 protein by Western blot (fibronectin 1, Supplementary Fig. 7a). In addition, TGF-β1 exposure resulted in a 0.25 log_2_FC reduction in *Survivin (Birc5)* expression after 12 hours in *Smad4*^Δ/Δ^ compared to a 2.55 log_2_FC reduction in *Smad4*^+/+^ adenoma. Anti-survivin antibody western blots and nuclear localisation suggested detectable reduction in protein levels in *Smad4*^+/+^ adenoma by 18 hours postTGFβ1 (Supplementary Fig. 8). The *Wnt* dependent intestinal stem cell marker *Lgr5* was downregulated in *Smad4^+/+^* following TGF-β1 (12 hours) compared to *Smad4*^Δ/Δ^ adenoma (2 log_2_FC, Supplementary Table 1), suggesting modification in Wnt pathway activity. Ecadherin and β-catenin proteins remained immunolocalised at the adherens junctions and in nuclei in both adenoma genotypes as expected (Supplementary Fig. 9). Immunoblots of adenomas from both genotypes showed multiple molecular weight bands for β-catenin and E-cadherin in the context of disruption of the β-catenin destruction complex by LOF of *Apc*^Δ/Δ^ (Supplementary Fig. 9b, c). Gene expression for *Myc* was not markedly different between genotypes, yet c-Myc protein levels appeared increased in *Smad4^Δ/Δ^* (12-18 hours) compared to *Smad4^+/+^*adenomas (Supplementary Fig. 9c, Supplementary Table 1). Overall, there did not appear to be significant correlation between changes in gene expression and protein localisation and abundance between genotypes. This implies that protein abundance, presumably regulated by variable processes independent of gene expression, is not a universal and reliable surrogate for gene expression in this context.

### Single cell RNA-seq of Smad4 ^fl/fl^ caecal adenoma are enriched in Pak3 ^Hi^ epithelial progenitor cells

In view of the differential changes in bulk mRNA expression, we next sought to quantify mRNA expression at the single cell level in the large adenoma directly harvested from the caecum. Epithelial cells (Epcam cluster) consisted of the *Lgr5* stem cell (*Lgr5* ISC) and seven stem cell-like (SC-L) clusters based on differential expression of *Lgr5* and other stem-cell markers such as *Lrig1, Smoc2, Hopx* as previously defined ^30–33^ (Fig. 4a, b. Supplementary Fig. 10, Supplementary Table 4). Differential enrichment in stem cell clusters was observed between the two genotypes, consisting of 61.8% of the total cell population in *Smad4*^fl/fl^ adenoma compared to 47.3% in *Smad4*^+/+^ (Figure 4c, d, Supplementary Table 3). A *Pak3*^High^ cluster (*Pak3*; P21 Rac1/CDC42 activated kinase 3) was the most enriched in *Smad4*^fl/fl^ (10.5%) compared to *Smad4*^+/+^ (2.5%) control adenoma. Pathway enrichment analysis for DEGs between *Smad4*^fl/fl^ and *Smad4*^+/+^ for the *Pak3*^High^ cluster showed an enrichment for the Wnt pathway and pluripotency markers (Supplementary Fig. 11). The same comparison in adenoma organoids between genotypes and all timepoints post TGFβ1 showed a non-significant difference in expression of *Pak3*, suggesting this was an *in vivo* adaptation that was not reproduced in organoids (Supplementary Table 1). The *Slc14a1^High^*cluster was also enriched in *Smad4*^fl/fl^ (7%) compared to *Smad4*^+/+^ (3.5%), where expression of the stem cell marker *Ly6a* was enriched (*Sac1*). Conversely, the *Lgr5* ISC cluster was significantly enriched in *Smad4*^+/+^ (5.8%) compared to *Smad4*^fl/fl^ adenoma (2.7%) (Figure 4d). *Smad4*^fl/fl^ showed significantly lower proportions of enterocytes (EC) as a consequence (7.8% *Smad4*^fl/fl^ vs 23.4% *Smad4*^+/+^) yet *Fmnl2^Hgh^*, *Cachd1^High^* and *Slc12a2^High^* showed similar proportions between genotypes. Note that the SC-L and the Lgr5 ISC clusters originate from the same node when viewed with cluster tree (0.2 resolution) (Fig. 4a). The *Tacstd2^Hgh^* cluster (murine foetal intestinal progenitor gene ^34^) and the *Cl^High^* cluster (*Clusterin* marker of the revival stem cell ^35^) are SC-L clusters with a similar proportions and originate from the same node (0 at 0.2 resolution). Markers for non-stem cells that lacked stem cell expression markers, included ribosomal gene expression ^33^ in transient amplifying cells(TA) and the Paneth cells marker *Pnliprp1* (pancreatic lipase-related protein 1) ^30^, enteroendocrine precursor *Gadd45g* ^33^, *Mt3* and *Pnliprp2* in secretory precursor cells (secretory pr).

**Figure 4.**
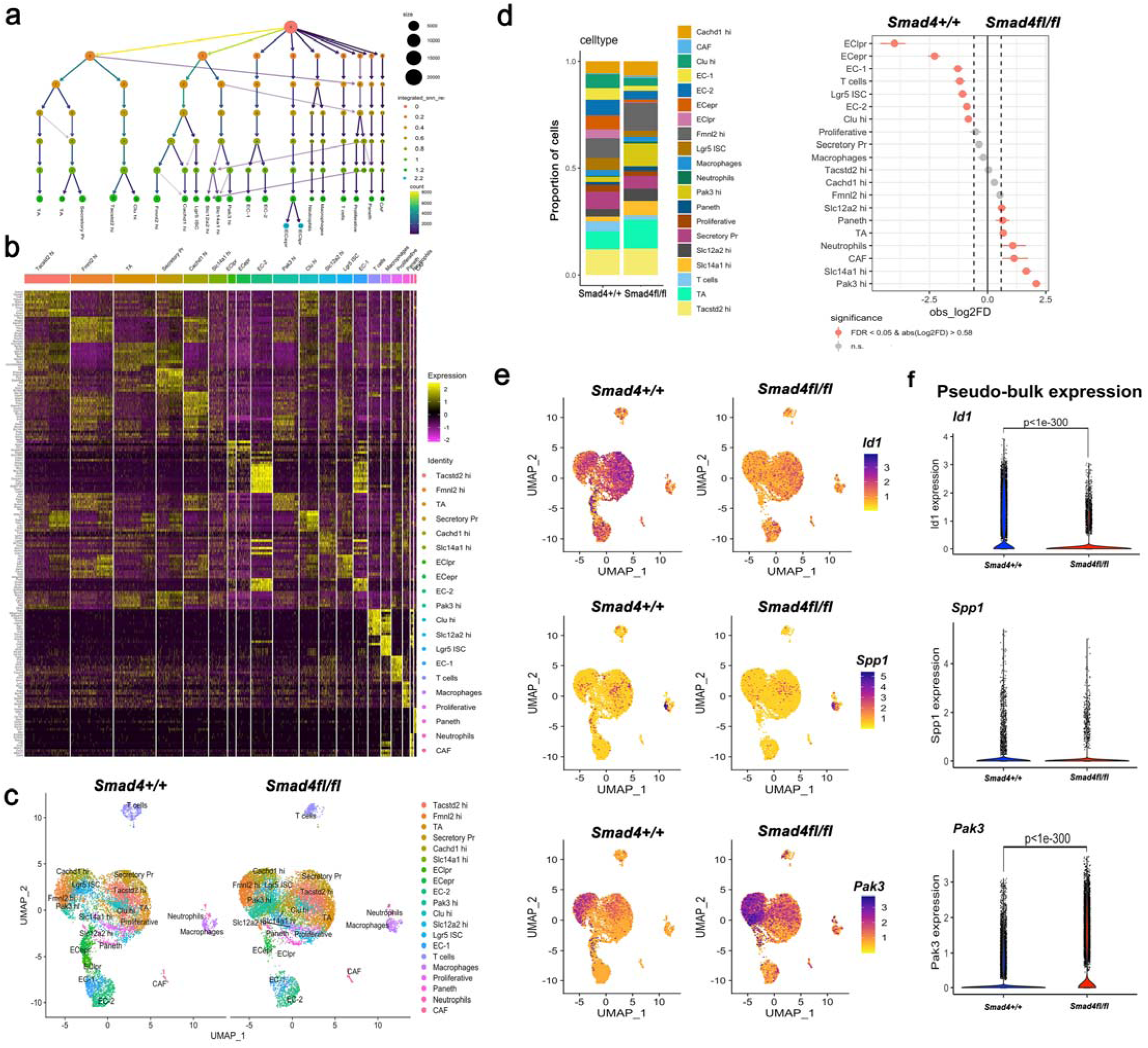
Smad4 dependent adenoma cell populations using single cell RNA-seq from in vivo caecal adenoma. a. Pooled caecum adenoma cells derived from different mice, *Apc*^fl/fl^*Smad4*^+/+^ n=3 and *Apc*^fl/fl^*Smad4*^fl/fl^ n=5, were subject to scRNA-Seq. Cluster trees show the resolution from 0 to 1.2 with the EClpr and ECepr clusters identified at resolution 2.2. b. Heatmap of the scaled expression of the top 5 genes identified in each cluster (genes selected based on the biggest difference between the cluster compared to all clusters). Lgr5 ICS = Lgr5 Intestinal stem cell; EC =enterocyte; EClpr =enterocyte late precursor; ECepr =enterocyte early precursor; TA= transit amplifying; CAF =cancer associated fibroblast; Secretory Pr =secretory precursor; hi =high referring to specific genes. c. UMAP plot of the cell-type clusters from *Apc*^fl/fl^*Smad4*^+/+^ (9674, n=3) and *Apc*^fl/fl^*Smad4*^fl/fl^ (11212, n=5) based on the expression of known marker genes. d. Bar graph; frequencies of clusters of the single cells in *Apc*^fl/fl^*Smad4*^+/+^ and *Apc*^fl/fl^*Smad4*^fl/fl^ adenoma. Point range plot; the confidence interval for the absolute log2FC (obs_log2FC) for the different identified cell types. Significant results are labelled in red. Note enrichment of *Pak3*^high^, *Slc14a1*^high^, CAF and neutrophil populations and the reduction in ECepr and EClpr populations in *Apc*^fl/fl^*Smad4*^fl/fl^ adenoma. Permutation test with FDR adjustment for p-value. e. UMAP plots comparing genotypes for single cell expression of *ID1* (suppression in all cell types in *Apc*^fl/fl^*Smad4*^fl/fl^), *Spp1* (monocytes and macrophage expression) and *Pak3* (increased expression in progenitor enterocyte population of *Apc*^fl/fl^*Smad4*^fl/fl^). f. Violin plots of pseudobulk RNA expression of *Id1*, *Spp1* and *Pak3.* Statistical analysis using MAST with Bonferroni adjustment for p-value.

Using both the 1.2 and 2.2 resolution cluster tree, we identified two groups of cells; the enterocyte early precursor (ECepr) expressing *Apoa1 Dmbt1, Reg4,* and *Reg3g,* and the enterocyte late precursor (EClpr) expressing *Reg3b, Mt1,* and *Mt2* ^33^. In *Smad4*^fl/fl^, EClpr (0.26% vs 4.1%) and ECepr (1.3% vs 6.5%) were depleted compared to *Smad4*^+/+^ adenoma. The mature enterocyte clusters, EC-1 and EC-2, showed expression of *Krt19, Krt8, Lgals3, Lypd8, Emp1,* and *Krt20* with differences in *Mxd1, Cdkn1a* and *Fam3*b expression. Significantly, 2.2% of *Smad4*^fl/fl^ were EC-1 cells compared to 5.4% for *Smad*^+/+^ (Fig. 4a, b, c, d). Thus, scRNA-seq of *in vivo* adenomas identified *Pak3^High^* and *Slc14a1^High^*as genes with increased expression and relatively differentially enriched in progenitor epithelial cells in *Smad4*^fl/fl^ caecal adenoma. Moreover, comparison of combined scRNAseq using a pseudobulk approach confirmed significant downregulation of *Id1* and upregulation of *Pak3* (Fig. 5f). In addition, *Id1* and *Sema5a* were the uniquely deregulated genes that were common between organoid bulk RNA-seq and the primary pseudo-bulk caecal adenoma scRNA-seq data (Supplementary Fig. 12). For *Spp1*, the high overall mRNA expression in adenoma derived organoids was not matched in the caecal derived single cell mRNA in epithelial cells, as expression was confined mainly to the monocyte-macrophage lineage (Fig. 6e).

**Figure 5.**
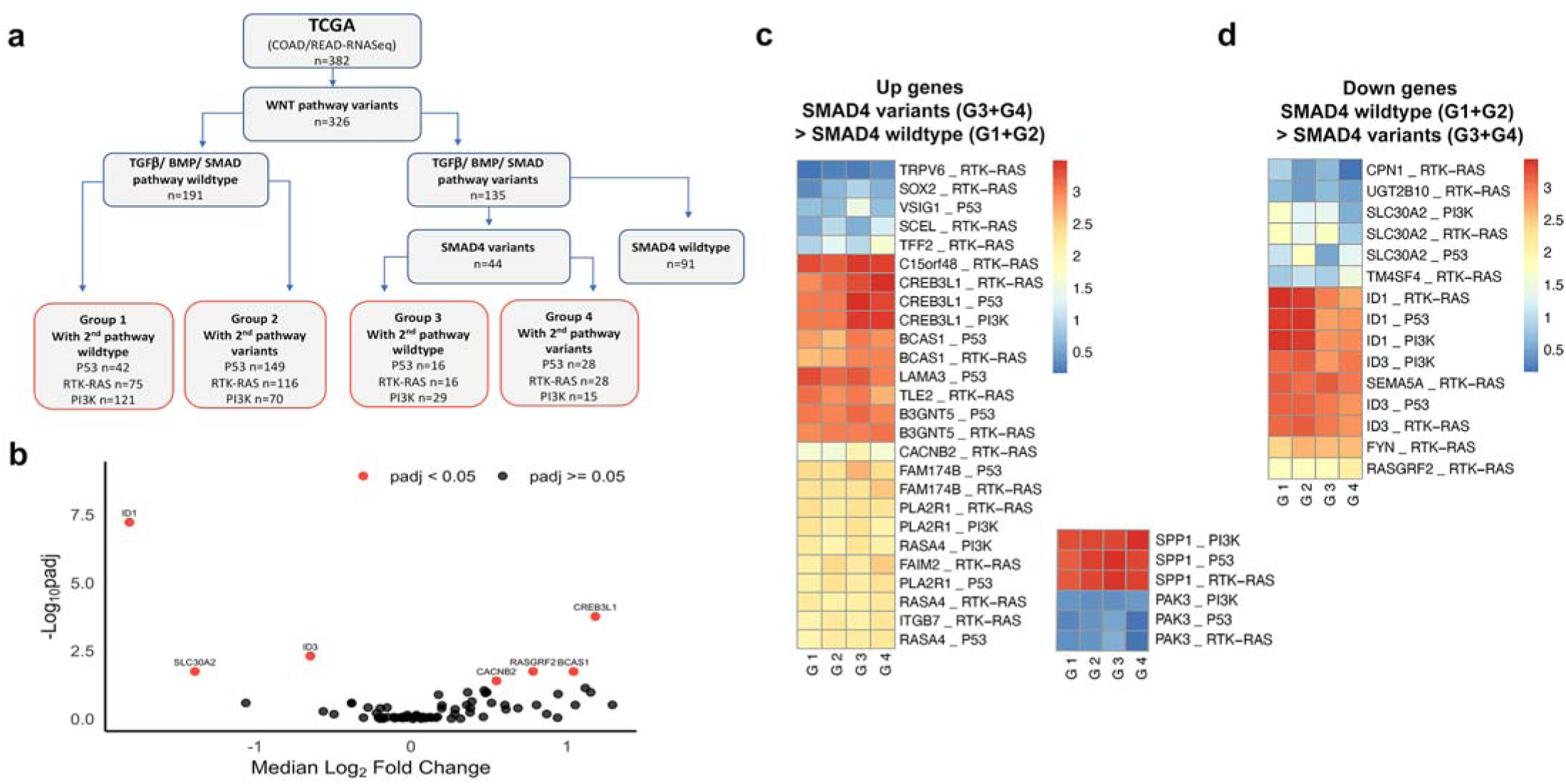
SMAD4 pathogenic variant dependent expression biomarkers for the Smad4-TGFβ gene-pathway in human colorectal cancer (TCGA) a. Flow diagram of TCGA bulk somatic DNA and RNA NGS sequencing data (COAD =colon adenocarcinoma, READ =rectal adenocarcinoma) with known WNT pathway driver variants (n=326). Pathogenic variants for TGF-β, activin or BMP receptors and Smad 2/3/4/5/7/9 were identified (n=135) from wild-type cases (n=191). Subgroups (G1-4) were then assembled with or without co-existing common pathway pathogenic variants in either RTK-RAS, TP53 or PI3K pathways. b. The human equivalents of the 76 significantly differentially expressed genes obtained from the comparison of bulk mRNA expression in the mouse adenoma were then compared in the human TCGA mRNA data set. Volcano plot of median log_2_ fold-change against adjusted p-value (-Log10) using Mann-Whitney U test (non-normal distribution using Shapiro-Wilk test) identified 7 human genes (in red) significantly different in expression between Smad4 wild-type (n=191) and Smad4 pathogenic variants (n=44)[BCAS1, CACNB2, CREB3L1, ID1, ID3, RASGRF2 and SLC30A2]. c., d. Comparison between sub-groups with and without context dependent variants G1G4 are shown as heat maps only for the significantly differentially expressed genes (median RSEM expression log_10_). Non-parametric Kruskal-Wallis test (FDR adjustment p<0.05) was followed by Dunn’s test and Holm adjustment for multiple comparisons. 26 gene expression-context pairs were identified to be upregulated in the TCGA cohort (c) and 15 gene expression-context pairs were significantly down regulated (d). *Spp1* and *Pak3* median expression are shown separately but were nonsignificant between groups (right heatmap). Of the comparison of the increased expression of *CREB3L1 and BCAS1* and repression of *ID1* and *ID3,* only *ID1* met the significance criteria.

**Figure 6.**
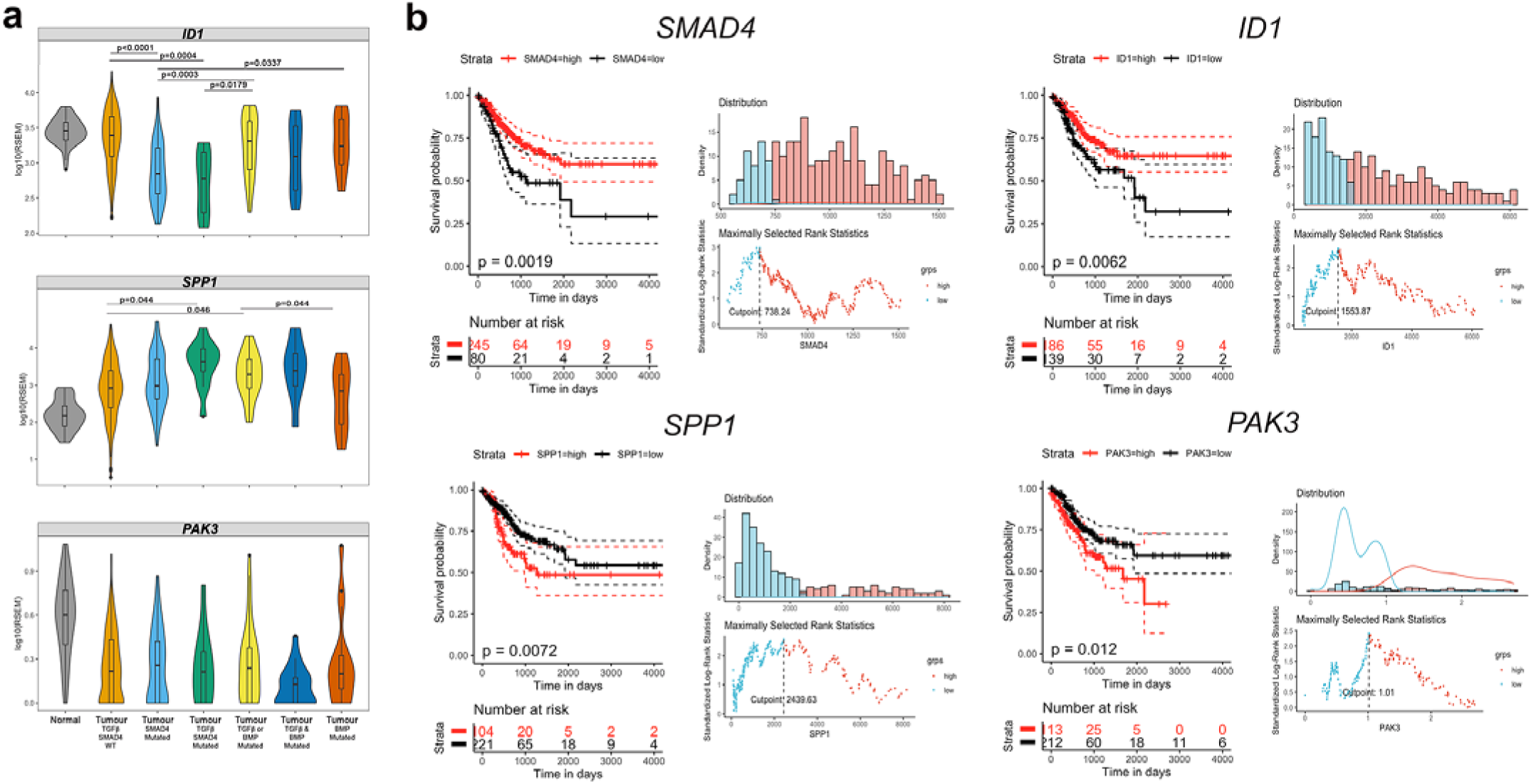
SMAD4, ID1, SPP1 and PAK3 expression biomarkers for the Smad4-TGF-β genepathway and human colorectal cancer (TCGA) survival. a. Comparison of *ID1, SPP1 and PAK3 mRNA* expression in cases with somatic pathogenic variants in either *SMAD4* alone, *SMAD4* with other genes in the TGF-β/BMP pathway (TGF-β *SMAD4* mutated), single gene variants in either the TGF-β or BMP pathway (TGF-β or BMP mutated), more than one variant in TGF-β /BMP (TGF-β and BMP mutated) or variants of BMP pathway genes alone (BMP mutated). Statistical analysis was performed using Kruskal-Wallis test (p <0.0001) followed by Dunn test multiple comparison test adjusted by Holm test (shown in the plot). Note the significant repression of *ID1* and increase in *SPP1* in *SMAD4* mutated cases. b. Kaplan-Meier survival curves showing survival differences between high and low gene expression of *SMAD4, ID1*, *SPP1* and *PAK3* (left). Cut points were determined using the maximally selected rank statistics (maxstat) method and are shown for each gene (right). Statistical significance between the High vs low expression groups determined with the Log-rank test (p values shown).

### Smad4 dependent ID1, SPP1 and PAK3 expression in human colorectal cancer

Having identified Smad4TGF-β1 pathway dependency of *Id1^low^*and *Spp1^high^* gene expression and *Pak3^high^* expressing cell enrichment in primary caecal intestinal adenoma, we next considered whether these functional gene expression biomarkers have prognostic relevance in human colorectal cancer (Fig. 5, Fig. 6, Supplementary Fig. 1, Supplementary Table 5). Data was derived from 382 individual cases from the human Cancer Genome Atlas project (TCGA; COAD, Colon Adenocarcinoma, READ, Rectal adenocarcinoma) that had both mutational profiles in c-BioPortal and RNA-seq data. We first tested whether the coexisting mutational context in colorectal cancer altered biomarker expression and key regulatory gene expression pathways by assembling mutational subgroups of colorectal cancer based on WNT-APC-β-catenin, RAS-MAPK, PI3K, and TP53 pathway variants (Fig. 5a). This test matrix derived four variant based sub-groups named Group 1-4 (Fig. 5a). Firstly, colorectal cancers with confirmed WNT-APC-β-catenin variants were selected and subsequently sub-divided. Group 1 and Group 2 cases were *SMAD4* wild-type and Group 3 and 4 cases had *SMAD4* pathogenic variants. Groups 1&3, and 2&4 differed depending on the presence or absence of second pathway variants in either the RAS-MAPK, PI3K or TP53 pathway, respectively. We next utilised the 76 DEGs identified in murine adenoma organoids and applied the equivalent genes to the human colorectal RNA-seq data. Volcano plots revealed only 7 genes that were significantly differentially expressed based on the presence (n=44) or absence (n=191) of a SMAD4 pathogenic variant (Fig. 5b). These genes were *BCAS1*, *CACNB2*, *CREB3L1*, *ID1*, *ID3*, *RASGRF2* and *SLC30A2*. Further comparison of the subgroups based on these genes with second pathway variants revealed 26 gene-context pairs that were significantly upregulated and 15 gene-context pairs that were significantly downregulated (Fig. 5c, 5d). Despite the correlation established for *Spp1* and *Pak3* in adenoma organoids, these two genes were not significantly differentially expressed in human colorectal cancer (Fig. 5c). Importantly, the upregulated expression of *CREB3L1 and BCAS1* and downregulated expression of *ID1* and *ID3* appeared least impacted by the co-existing 2^nd^ pathway variants based on visualisation of the heat maps. However, analysis based on our three predefine criteria (see Methods) of the comparison of the increased expression of *CREB3L1* and *BCAS1*, and repression of *ID1* and *ID3*, only *ID1^low^* met the significance criteria. Analysis of the TCGA cohort with respect to either the presence or absence of any additional *TGF-β/BMP* pathway variants also showed that *ID1* expression was also significantly downregulated with co-existing *SMAD4* and other *TGF-*β pathway downstream mutations compared to normal control tissues (Supplementary Table 5). Compared to non-SMAD4 associated TGF-β/BMP pathway pathogenic variants (e.g. in Type 1 receptors, other SMADs or in the BMP pathway), *SMAD4* mutations specifically appeared to have the greatest impact on overall *ID1* repression (Fig. 6a, p<0.0001). In the context of all combined TGF-β/BMP pathway mutations, however, *SPP1* mRNA did appear to be expressed significantly higher compared to cases without mutations in the TGF-β and BMP pathway (p=0.044, respectively, Dunn test, Fig. 6a). For *PAK3*, no significant overall expression differences were observed between *SMAD4* mutational groups (Fig. 6b). We next evaluated whether any of the expression biomarkers impacted on the survival outcome of colorectal cancer using maximum selected rank statistics to define a single cut-point (Fig. 6b). For example, *Smad4^low^* expression (p=0.0019), *ID1^low^* expression (p=0.0062), *SPP1^Hi^*^gh^ (p= 0.0072) and *PAK3^High^* (p=0.012) appeared to be associated with poorer overall survival in the TCGA cohort (Kaplan-Meier, Fig. 6b). The less significant impact of *PAK3^High^* may reflect the relative low proportion of *PAK3^High^* expressing sub-populations of cells within tumours.

## Discussion

The identification of cancer cells that have activation of the TGF-β-BMP-SMAD4 pathway in the context of pathway gene disruption requires validated models in order to discover mechanism and pathway specific gene expression biomarkers ^2^. Moreover, RNA based molecular prognostic classifiers of colorectal cancer have developed in line with the now clinically applied breast cancer equivalents. Such machine learning classifiers have identified several clusters (e.g. CRISB) that include a stromal based signature that correlates with poor prognosis in colorectal cancer ^3,36,37^. A supervised machine learning CMS1-4 classifier (Consensus Molecular Classifier) has correlated gene level expression signatures with four prognostic groups, where the poor prognostic group CSM4 was associated with EMT and TGF-β dependency, yet the classifier does not utilise RNA expression biomarkers that are specific readouts of the TGF-β-BMP-SMAD4 pathway. With respect to the latter, human colorectal polyp derived organoids have been recently used to discover TGF-β dependent gene sets in human serrated adenomas combined with a *BRAF^V600E^* mutations, where there was an induced mesenchymal phenotype, yet this model did not address specific *Smad4* loss of function ^38^. Further analysis of human expression data has recently identified three classes of pathway derived subtypes (PDS1-3) in colorectal cancer with and without *KRAS* mutations, where CSM4 and CSM1 combine in the PDS2 subtype where TGFβ-BMP-SMAD4 pathway expression profiles appear enriched ^4^. These subtypes appeared independent of *KRAS* driver gene variant classes and phenotypic appearance and for PDS1 and 2, correlated with RNA expression data from mouse genetically modified models. Thus, gene-pathway specific gene expression biomarkers their dependencies are overall becoming better understood and applied in intestinal cancer classifiers.

In a murine system, we discovered that the generation of homozygous *Smad4* with *Apc* LOF alleles was associated with a discordant intestinal phenotype, with adenoma suppression in the small intestine and adenoma progression in the caecum. By deriving organoid cultures from the adenoma, we compared control wild-type *Smad4* (*Smad*^+/+^) with *Smad4* LOF (*Smad4*^Δ/Δ^). We discovered that the high sensitivity to TGF-β1 induces similar EMT and cell death phenotypes in a dose dependent manner, with a relative 10-fold dose resistance to TGF-β1 in *Smad4*^Δ/Δ^ adenoma. Following TGF-β1 exposure, we identified genes with extremes of differential expression by genotype, including reduced *Id1* and increased *Spp1* gene expression as consistent features in *Smad4*^Δ/Δ^ murine adenoma organoids. The gene expression changes as consistent with *Smad4* independent activity of the RSmads, in this case this is likely to be mainly Smad3 ^39–41^. Single cell RNA sequencing of the primary caecal adenomas detected enrichment of the *Pak3^High^* expressing progenitor epithelial cell population in *Smad4^Δ/Δ^*. We assume that that the differentially expressed genes are the main underlying difference that may account for functional discordant progression of adenoma, for example *Id1^Low^*.

Mouse models of *Smad4* LOF in the intestine have been validated and have shown that *Smad4*^+/^mice spontaneously developed juvenile polyposis (JP) at 1.5 years, and invasive adenocarcinoma when crossed with *Apc*^Δ7^^16/+^ ^42-45^. In addition, an *Apc*^1638N/+^*Smad4*^fl/fl^*K19-CreERT2* model showed an increased adenoma burden^46^, and CRISPR/Cas9-mediated genome editing of human intestinal epithelial organoid cultures introduced mutations in *APC*, *TP53*, *KRAS* and *SMAD4* which, when combined and selected, developed aggressive invasive carcinomas^23^. Moreover, *SMAD4* LOF blocked differentiation during tumour progression in an orthotopic organoid transplantation model mimicking an intestinal adenoma-carcinoma transition ^47^. These results may also help to explain the selection of deletions of chromosome arms 18p and 18q (including *SMAD4*) detected in 66% of invasive colorectal cancers ^48^.

The generation of homozygous alleles in our conditional model results in rapid adenoma formation within days, with animals reaching humane endpoints because of anaemia and intestinal obstruction before the detection of invasive and metastatic sites. The discordant pattern of adenoma development in *Apc*^fl/fl^*Smad4*^fl/fl^ suggests that the colon may be more prone to adenoma initiation and progression when both genes are disrupted. One possibility is that this may be due to the caecal inflammatory microenvironment and the local TGF-β-BMP ligand supply. Our RNA-Seq analysis showed downregulation of *Pla2g2a* (2.5log_2_FC) in *Smad4*^Δ/Δ^ adenoma. *Pla2g2a* is considered to be a tumour suppressor independent of the Wnt/β-catenin signalling pathway, and intraluminal secretion of Pla2g2a is a host defence mechanism during the active phase of ulcerative colitis ^49^. Moreover, Reg3y and *Reg3*β were also downregulated (1.2log_2_FC) in *Smad4^Δ/Δ^* adenomas and as both genes are important for epithelial defence against bacterial invasion ^50,51^. Another confounder may be modifier alleles co-segregating with LoxP alleles, although this was mitigated by backcrossing to >10 generations of all conditional alleles to C57Bl/6J ^27^. Excess TGF-β-BMP may also promote adenoma formation independent of *Apc* LOF, as targeted deletion of *Bmpr1a* can lead to crypt expansion, fused villi and adenoma formation due to an increase in β-catenin ^52^. Overall, adenoma *Smad4* LOF may create the conditions for local TGF-β1 ligand supply and further stimulation of the pathway, contributing to differentiation arrest, chronic inflammation and subsequent adenoma progression.

The dose dependency of TGF-β1 ligand supply with respect to cell growth phenotype has not been previously described to our knowledge, but may contribute functionally to the manifestation of differential phenotypes with *Smad4* LOF; at a given dose, pathway gene expression positively modifies differentiation and EMT function, but at the same time there is resistance to cell death and cell cycle arrest. The time-dependent TGF-β1 on gene expression, using a fixed concentration of 390pM which exceeds the IC_50_ of *Smad4^+/+^*adenoma, resulted in differential expression of genes that were *Smad4* and TGF-β1 dependent, with the most significant being decreased expression of *Id1. Id1* codes for a non-DNA binding inhibitor of basic-helix-loop-helix (bHLH) transcription factors (e.g. GATA4) to inhibit multiple cell processes including differentiation. ChiP-Seq assays that have shown that Smad3 and Smad4 can bind the *Id1* promoter to directly stimulate early induction following TGF-β1 exposure ^40^. Feedback *Id1* repression is then thought to be also pSmad3 dependent but through feedback activation of *ATF3*, that then directly represses *Id1* expression ^53^*. Id1* over-expression in pancreatic ductal adenocarcinoma in mouse and human is also associated with more invasive and undifferentiated cells, a mechanism proposed to be dependent on EIF2 signalling ^54^ and independent of the TGF-β pathway^55^. Id1 proteins detected by IHC show increased expression with in colorectal cancer and mouse models, and one report (although the paper was subsequently retracted) showed that a genetic knock-out of *Id1* results in adenoma suppression in the small intestine of *Apc^Min/+^*but not when colitis is chemically induced, in support of an inflammatory basis of the caecal adenoma ^56^. Id1 protein localisation has not been shown to correlate with gene expression, as we have observed, and so is unlikely to be a reliable functional biomarker of the pathway activity ^56^. Recently, a Nur77 based molecular switch has also been proposed to regulate *Id1*, where stabilisation of TGF-β induced Smad3 degradation leads to an increase in *Id1* expression, whereas in the absence of TGF-β, Nur77 enhances Smad3 degradation and results in reduces *Id1* expression^57^.

*Spp1* (osteopontin) expression normally occurs in osteoclasts and macrophages, but has also been specifically detected in tumour associated M2 macrophages, with senescence associated secretary phenotype (SASP) epithelial tumour cells and at the invasive front of human colorectal tumours ^58–62^. As well as extensive circumstantial data associating high *SPP1* expression in human colorectal cancer, *Spp1* was also previously reported to be a key *Smad4* dependent expressed gene functionally associated with murine prostatic cancer invasion in a *Pten* prostate deletion model ^63^. High *SPP1* expression may have therapeutic implications, as immune exclusion correlates in the tumour microenvironment with cancer associated fibroblasts resulting in resistance to immune checkpoint therapy^64,65^.

*Pak3* is a well-known downstream effector of Cdc42 and Rac1 GTPases that mediates motility and MAPK activation associated with cancer invasion and metastasis ^66–68^. In a recent adenovirus Cre-induced *Kras^G12D^, p53*^fl/fl,^ *Smad4*^fl/fl^ model of lung cancer, increased metastasis was associated with *Smad4* LOF and *Pak3* activation, the latter identified as a downstream effector of *Smad4* via the PAK3-JNK-Jun pathway ^68^. Interestingly, the proposed mechanism appeared to be due to attenuation of *Smad4* dependent transcription of miR-495 and miR-543 that can directly bind to the PAK3 3’UTR. Overexpression of PAK family members are also associated with CRC outcome, transition of adenoma to carcinoma and stabilisation of β-catenin ^69–75^.

Finally, having determined 76 differentially expressed TGF-β-BMP-SMAD4 genepathway dependent candidate gene expression biomarkers in mouse, we evaluated whether these same genes in humans were also differentially expressed with respect to the presence or absence of SMAD4 pathogenic variants. At least 7 such DEGs were identified, several of which (*CREB3L1, BCAS1, ID1* and *ID 3*) also appeared independent of common co-existing pathogenic variants in the RTK-RAS, TP53 and PI3K pathways. *SMAD4*, *ID1, SPP1* and *PAK3* mRNA expression also have prognostic implications for colorectal cancer using the TCGA cohorts, and are potential functional readouts of the TGF-β-BMPSMAD4 pathway. We provide evidence these genes, and potentially their associated pathway gene sets, should be incorporated into future prospective evaluation of pathwaybased prognostic-predictive classifiers for the TGF-β-BMP-SMAD4 pathway in intestinal cancer subtypes^4^.

## Materials and methods

### Conditional Cre intestinal adenoma model

Work was performed under a UK Home Office licence, and approved by University of Oxford Animal ethics committee. The mouse lines were backcrossed to C57Bl/6J as reported including *Smad4*^fl/fl^ (*Smad4*^tm1Rob^) (Supplementary Material and Methods). Genotyping PCRs were briefly (5’-3’, forward-F and reverse-R): *Apc* FGTTCTGTATCATGGAAAGATAGGTGGTC and RCACTCAAAACGCTTTTGAGGGTTG (GAG-TACGGGGTCTCTGTCTCAGTGAA for recombined product), *Smad4* FCTTTTATTTTCAGATTCAGGGGTTC (F-AAAATGGGAAAACCAACGAG for recombined product) and RTACAAGTGTATGTCTTCAGCG. Genotype combinations were generated by experimental breeding. *Smad4*^fl/fl^*RosaYFP*^fl/fl^ and *Apc*^fl/fl^*Smad4*^fl/fl^*RosaYFP*^fl/fl^ were crossed with either *Lgr5*-*CreER*^T2^ or *Vil*-*CreER*^T2^ mice. *Apc*^fl/fl^*Smad4*^+/+^*RosaYFP*^fl/fl^ *Lgr5*-*CreER*^T2^/*Vil*-*CreER*^T2^ animals were used as controls. Mice bred with *Vil*-*CreER*^T2^ mice were used for an *in vitro* experiment only. For *Lgr5-CreER^T2^*activation, adult mice age ≥ 8 weeks had an intraperitoneal injection of 5mg (200 mg.kg^-1^) tamoxifen dissolved in corn oil. Mice were monitored daily, small intestinal and colonic adenomas were scored for number, diameter, distribution and weight as described^27^.

### Intestinal adenoma organoids

Adenoma formed *in vivo* from *Smad*^fl/fl^ or *Smad4*^+/+^ Lgr5 mice were cultured as described^27,76^. For crypt culture, the small intestines were collected from *Smad4*^fl/fl^*RosaYFP*^fl/fl^ and *Apc*^fl/fl^*Smad4*^fl/fl^*RosaYFP*^fl/^ *Vil*-*CreER*^T2^ mice and processed as described ^76^. Briefly, crypts were cultured in Matrigel and supplemented with ADF2 and crypt growth factors, ENR (8.1 nM EGF (E), 500ng ml-1 R-spondin1 (R), and 2.16nM Noggin (N)) and incubated at 37°C, 5% as described. At D3, crypts generated from transgenic mice were incubated with 500nM 4-hydroxy-tamoxifen (4-OHT) (Millipore, 579002) dissolved in 100% EtOH (test), vehicle (negative control) with 5µl EtOH for 12h at 37°C. The test and the vehicle treated organoids were supplemented with EN only (no R) and the normal crypts with ENR. Proteins were extracted from organoids as described ^77^.

### Cell labelling and imaging

Neutral buffered formalin fixed tissues were paraffin embedded and 5µm sections were processed either by immunohistochemistry (IHC) or immunofluorescence (IF). The antibodies that used in this study are listed (Supplementary Materials and Methods). Organoids growing on coverslips in 24 well plates were used for IF. Organoid proliferation used the thymidine analogue 5-ethynyl-2’-deoxyuridine (EdU) for 2 hrs using Click-iT EdU Alexa Fluor-555 Imaging Kit (Invitrogen, C10338). H&E and IHC slides were viewed by brightfieldfluorescent microscopy (Olympus BX60). Images were acquired by CCD camera and Nuance 2.10 software (CRI, Massachusetts). Confocal images were captured with an Olympus FluoView FV1000 using 40x 1.3NA lens and associated software. Labelled slides (EdU, IHC or IF) were analysed using ImageJ 1.48 software. AlamarBlue (AB) (AbD Serotec, BUF012B), was used to quantify adenoma growth. Absorbance after 24hr was read at 570nm and 600nm using a Spectramax M5 plate reader (MTX Lab Systems). Live cell imaging was conducted with a Zeiss Axiovert 200 inverted microscope with a BSI Prime CMOS camera with a Neofluar 10x 0.3NA lensfor adenoma organoids treated with TGF-β for 24hrs. MetaMorph software (v 7.8.12.0) multidimensional Z-stack acquisition with 100 milliseconds exposure acquired at regular intervals of 15 minutes over an extended duration of 6-18 hours culture at 37.5 °C, and 5% CO_2_ in air.

### TGF-*β*1 induced *g*ene expression bulk RNASeq

Early passage adenomas seeded at 9,000 cells per 25µl well were used for RNA-Seq and miRNA-Seq. Adenomas were grown in ADF2 medium supplemented with 8.1nM EGF, 10µM Y-27632 and 1µM SB-505142. The medium was changed on D2, the cells were washed, EGF and 390pM TGF-β1 added and samples extracted from three different time points (t=0, t=1, t=12 hours post TGF-β1 treatment). Three biological replicates from each genotype and RNA was pooled from 8 wells per treatment. RNA extraction was performed using Quick-RNA MiniPrep (Zymo Research, R1054), assessed using a Nanodrop-1000 spectrophotometer and RNA Clean & concentrator^TM^-5 (Zymo Research, R1015). RNA-Seq and miRNA-Seq were performed at the Wellcome Trust Center for Human Genetics, University of Oxford. Briefly, directional multiplexed mRNA and miRNA Libraries were prepared. All mRNA was ribo-depleted and converted to cDNA. Paired-end sequencing (100nt) were run on a HiSeq 2500 (Illumina) for mRNA-Seq.

Transcript abundances from RNA-Seq data were quantified using the Tuxedo pipeline^78^. Fastq reads were aligned to the mouse genome (mm10/GRCm38) with the TopHatBowtie2 aligner, versions 2.1.0 and 2.3.4, respectively and expression of transcripts was quantified with Cufflinks 2.2.1, as fragments per kilobase per million mapped reads (FPKM). Counts were also generated using the feature ‘Counts’ from the Subread package. Pairwise differential expression analysis between groups was performed in R using DESeq2 version 1.28^79^. Genes that have less than 5 reads in at least 3 samples were removed. Genes with adjusted p-values (FDR) < 0.05 were considered significant. Shrunk log2fold change (log2FC) was calculated using as a shrinkage estimator. Genes considered as differentially regulated between 2 conditions have ±1.5log2FC. Time-series analysis was performed using the likelihood ratio test implemented in DESeq2. Gene ontology term enrichment analysis for the significant DEG genes performed using “clusterProfiler” where we evaluated biological processes and molecular functions^80^. For easier visualisation, we used “tmod”package for enrichment analysis (https://peerj.com/preprints/2420v1). In tmod, we applied CERNOtest, Coincident Extreme Ranks in Numerical Observations (CERNO) after converting mouse genes to human orthologs. We performed the enrichment analysis using Hallmark gene sets from MSigDB v7.2 (Broad Institute) and the blood signature from LI modules in tmod, gene sets with the area under the curve (AUC) ≥ 0.5 and padj<0.05 were considered positive. Metascape website was utilised for pathway analysis of certain gene lists, and to identify the transcription regulatory network based on TRRUST ontology^81^.

Early TGF-β target genes were classified based on log2FC ±1.5 expression and adjusted p <0.05 after 1h compared to t=0 in *Smad4^+/+^*adenomas. Transient early upregulated/ downregulated genes were the genes with opposite directions to their initial regulation when compared to late time points (t=12 vs t=0 and/ or t=12 vs t=1 hour) in *Smad4^+/+^* adenomas. Late TGF-β upregulated/downregulated genes were classified according to their upregulation/downregulation at 12 hours in *Smad4^+/+^* adenomas compared to t=0 and/or t=1 hour with log2FC ±1.5 and adjusted p <0.05. Genes were considered *Smad4*independent if they were significantly expressed in both *Smad4*^+/+^ and *Smad4*^Δ/Δ^ adenomas with TGF-β1 treatment compared to their corresponding untreated baseline. However, if genes were differentially expressed in an opposite direction to that of TGF-β target genes in *Smad4^Δ/Δ^* vs *Smad4^+/+^* at t=0, they were classified as Smad4-dependent genes and excluded from the *Smad4^Δ/Δ^* vs *Smad4^+/+^* t=0 list. TGF-β target genes that were only upregulated in *Smad4^+/+^*adenomas were classified as *Smad4*-dependent genes. Genes that were expressed in *Smad4^Δ/Δ^* vs *Smad4^+/+^* at t=0 hour and were not among TGF-β target genes and considered for baseline comparisons.

### Caecal adenoma single cell mRNA sequencing

Pooled single-cell suspensions from were obtained from caecal tumours from three *Apc*^fl/fl^*,Smad4*^+/+^, *Lgr5Cre*^ERT2^ and five *Apc*^fl/fl^*, Smad4* ^fl/fl^*, Lgr5Cre* ^ERT2^ mice. Cell suspensions underwent library preparation using the Chromium Single Cell 3’ kit (10X Genomics) and sequencing using Illumina HiSeq 2×150bp (Azenta, USA), with approximately 25,000 reads per cell. Sample demultiplexing, barcode processing and single-cell 51 unique molecular identifier (UMI) counting (Cell Ranger Software Suite v7.0.1) with FASTQ alignment to the mouse reference transcriptome mm10-3.0.0. The following criteria were then applied to each cell: gene number between 250 and 7,000, UMI count >500 and mitochondrial gene percentage <0.15, log10 Genes Per UMI >0.80. Doublets were removed using the scDblFinder library. Genes with ten read counts or less were removed. A total of 20886 cells were analysed (11212 for *Smad4^fl/fl^* and 9674 for *Smad4^+/+^*). Dimensionality reduction and clustering used the filtered gene-barcode matrix normalised using the SCTransform with Seurat (v4.3.0) ^82^. Variables including mitochondria Ratio, nUMI, nGene and the cell cycle gene difference were regressed and the top 3000 variable genes were used. The filtered gene-barcode matrix of all samples were then integrated, k nearest neighbour (k-NN) generated using FindNeighbors (Seurat) using the top 40 principal components and clusters identified using FindCluster function (Louvain algorithm). We used resolution 1.2 to classify the different cell types and resolution 2.2 to determine the enterocyte precursors based on clustree (v0.5.1) ^83^. Gene markers that identified each cluster relative to all other clusters was determined with FindAllMarkers function in Seurat using the Wilcox test (log-fold change threshold 0.25). Manual cell labelling of the top marker genes for each cluster utilised previous scRNA-seq studies ^30,32,84^. Significant differences (p values, and confidence intervals via bootstrapping) in cell type proportion between each genotype was determined with the permutation test implemented in scProportionTest package in R (V4.2.3). Data was visualised using Uniform Manifold Approximation and Projection (UMAP) dimensional reduction and plots were generated using Seurat and the scCustomize library. Each cluster was determined using FindAllMarkers function in Seurat (Wilcox test). The log fold change threshold was set to 0.25. Manual cell labelling utilised gene clusters in previous studies ^30–33^, the Lgr5 intestinal stem cell signature ^32^ and the CellMarker 2.0 database ^84^. The significant differences in cell type proportion used the permutation test implemented in scProportionTest package in R which calculate a p-value for each cluster, and a confidence interval for the magnitude difference via bootstrapping ^85^. All analysis was performed in R V4.2.3. Differential gene expression analysis for each cluster between *Smad4*^fl/fl^ and *Smad4+/+* were performed using MAST ^86^ with FindAllMarkers function in Seurat. Differentially expressed genes were selected based on log fold change threshold and expressed in at least in 25% in one of the groups. Pathway analysis was performed using Metascape ^81^. Pseudo-bulk expression analysis was performed using MAST cell implemented in Seurat’s FindMarkers function with the first and second identities using all *Smad4+/+* and *Smad4*^fl/fl^ cells, respectively.

### TCGA colorectal cancer analysis

Normalised gene expression from RNA-seq (illumine hiseq Level 3 RSEM normalized genes) retrieved from http://firebrowse.org/?cohort=COADREAD accessed October 2019. Genes and pathways that are frequently mutated in colorectal cancer, retrieved from Cancer Genome Atlas project (TCGA)^48^, include the Wnt/β-catenin pathway (*APC, CTNNB1, DKK1, DKK2, DKK3, DKK4, LRP5, FZD10, FAM123B:AMER1, AXIN2, TCF7L2, FBXW7,* and *ARID1A*), TP53 pathway (*ATM, TP53*). PI3K (*IGF2, IRS2, PTEN, PIK3R1, and PIK3CA*), RTKRAS pathway (*NRAS, KRAS, BRAF, ERBB2, and ERBB3*). TGF-β pathway (*ACVR1B, ACVR2A, SMAD2, SMAD3, SMAD4, TGFBR1* and *TGFBR2*) and BMP pathway (*SMAD1, SMAD9, SMAD5, BMPR1A, BMPR1B* and *ACVR 1*). Genomic variants (missense mutation, homozygote deletion, amplification or fusion gene) in these genes for COAD and READ TCGA cohorts were retrieved from cBioPortal (https://www.cbioportal.org), accessed December 2019. In COAD and READ TCGA cohorts the variant status was available for 56 patients with *SMAD4* variants. 98.1% of these patients had alterations in the Wnt/β-catenin pathway, 59.3% had alteration in the TP53 pathway, 46.3% in the PI3K pathway and 64.8% had variants in the RTK-RAS pathway. Normalized RNA-seq was available for 51 normal tissues and 326 patient tumours with variant data. In TCGA COAD-READ cohorts, 44 patients with probable pathogenic and possibly pathogenic SMAD4 variants, but with no other variants in the TGF-β pathway, had RNA-seq data. One patient did not have a variant in the Wnt pathway and was excluded from the analysis. Patients with known variants in the TGF-β pathway were excluded from the control group. Using this patient group, we validated the top 76 DEG in the mouse adenomas, genes with ±2log2FC and padj<0.05 from Smad4^Δ/Δ^ vs Smad4^+/+^ (t=0 hr), either in the presence or absence of variants in the PI3K, TP53, PTK-RAS pathways in the context of *SMAD4* variant. Statistical analysis was performed using Kruskal-Wallis test followed by FDR adjustment. Genes with p<0.05 following FDR adjustment were considered significant. Multiple comparisons were performed using Dunn test followed by Holm adjustment to detect *SMAD4*-dependent genes. The expression between 2 groups were considered positive if adjusted p<0.05. We considered genes to be *SMAD4* dependent genes based on three criteria. First, there needs to be a significant difference in gene expression between the groups (Group 1-Group 3, and Group 1-Group 4, Group 2-Group 3 and Group 2-Group 4) in the context of the three second pathways with and without variants. Second, there should be no significant difference in gene expression between Group 1-Group 2, and between Group 3 -Group 4, respectively. Third, the any change in gene expression should follow the same direction as observed in the mouse adenomas. Note that 84 genes differentially regulated at t=0 with absolute log2FC >=2 and adjusted p<= 0.05. Of those, 76 genes had human orthologs, and 73 genes had RNA data in the TCGA cohorts. Survival data for the COAD-READ TCGA cohort were retrieved (Table S1, Tab TCGA-CDR, accessed May 2019^87^).

### Statistical analysis

Statistical and graphical data analyses were performed using either Graph-Pad Prism version 6.0. software or R version 4.0.2. Images were analysed using ImageJ 1.48 software. Final Figures were generated using Photoshop (CS5). Immunoblot analysis was performed as follow; loading control and test from scanned immunoblots was aligned horizontally on Photoshop (CS5) and analyzed using ImageJ 1.48 software. For TCGA COAD-READ expression data, the cut-off point between high and low mRNA expression levels for each gene was computed based on the maximally selected rank statistics (maxstat) using the surv_cutpoint function from the ‘survminer’ package in R.

## Acknowledgements

We thank Elisabeth Robertson for the conditional Smad4 model and Caroline Hill for discussion. Linda Randall and Nicky Hamp for mouse technical assistance.

## Author Contributions

Experiment conceived by ABH. Mouse breeding and phenotypic analysis by MS and JM, imaging by MS, SB, and CB, bulk gene expression MS and DB, genotyping by MS, EC, and JH, histopathology HM, evaluation of cancer cell model systems, MS, PM and CJI, single cell sequencing analysis MS, TCGA analysis, MS, manuscript was written by ABH, MS and DB. All authors approved the manuscript.

