## Supplementary Figures for "*Smad4* and TGF-β1 dependent gene expression biomarkers in conditional intestinal adenoma, organoids and colorectal cancer"

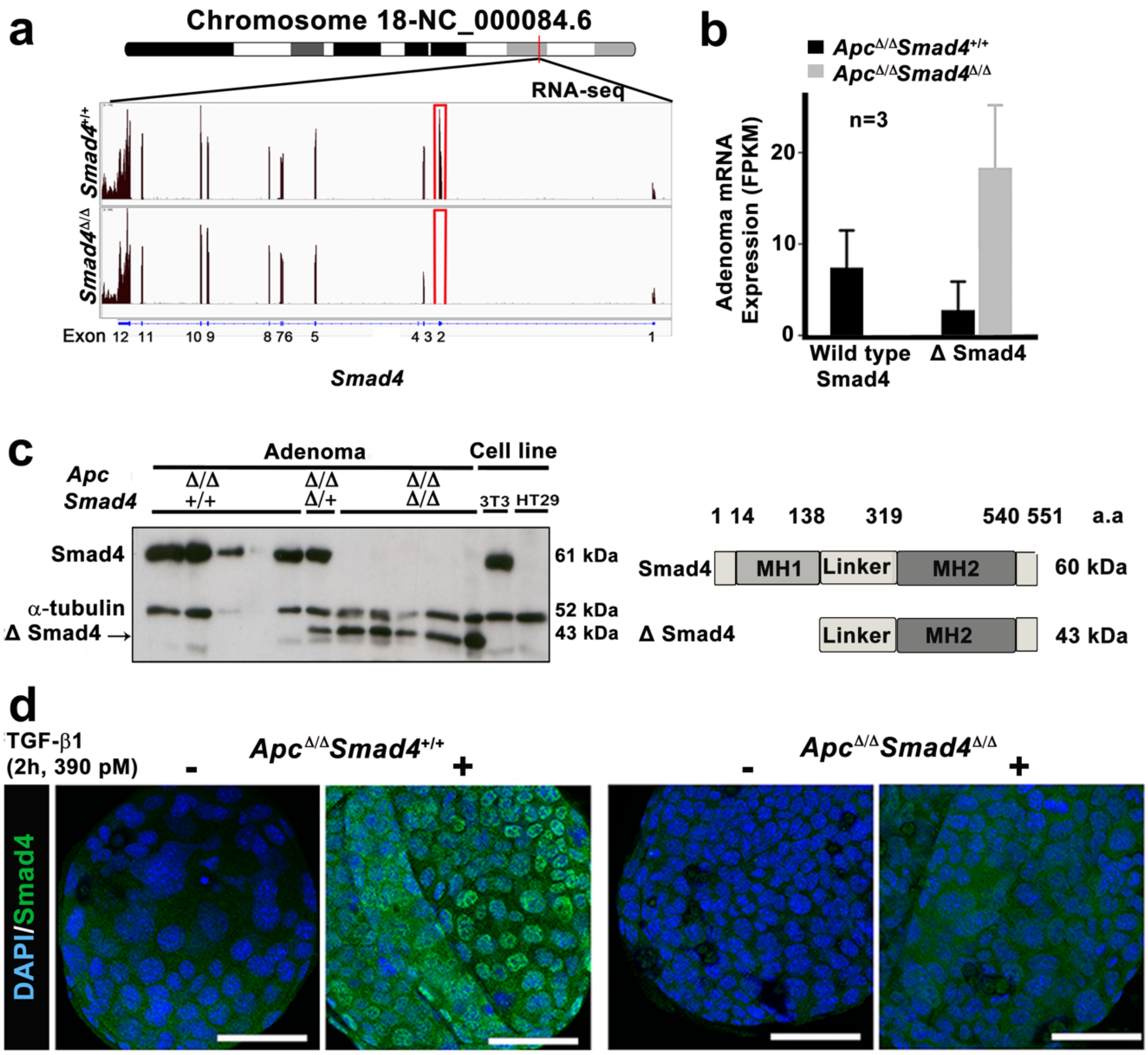


**Supplementary Figure 1**
**MH1 deletion in *Apc*$^{\Delta/\Delta}$*Smad4*$^{\Delta/\Delta}$ following Cre-mediated recombination.**

**a.** IGV coverage track for *Smad4* mRNA. Bam files generated from RNA-seq for the *Smad4* gene on Chromosome18; NC_000084.6 Locus. **b**. Quantification of mRNA abundance in FPKM derived from *Apc*$^{\Delta/\Delta}$*Smad4*$^{\Delta/\Delta}$ and *Apc*$^{\Delta/\Delta}$*Smad4*$^{+/+}$ adenoma organoids. **c.** Western blots (left panel) showing Smad4 protein of 61 kDa from *Apc*$^{\Delta/\Delta}$*Smad4*$^{+/+}$ and *Apc*$^{\Delta/\Delta}$*Smad4*$^{\Delta/+}$ adenomas, and a truncated Smad4 protein of 43 kDa in *Apc*$^{\Delta/\Delta}$*Smad4*$^{\Delta/+}$ and *Apc*$^{\Delta/\Delta}$*Smad4*$^{\Delta/\Delta}$. Controls: 3T3 fibroblast cells with intact *Smad4* (positive control). HT29 CRC cells have *Smad4* mutation (C931 C>T) in the MH2 domain (negative control). Simplified cartoon (right panel) representing mapping with ExPASy shows deletion of the MH1 domain leaving an intact linker and MH2 domain of Smad4, following *in silico* analysis that showed an open reading frame in exon 5 of *Smad4*. The predicted size of truncated Smad4 is 43 kDa. **d.** Representative confocal images for Smad4 nuclear localisation in cultured adenoma following 2hr of 390pM TGF-β1 treatment (n=4).

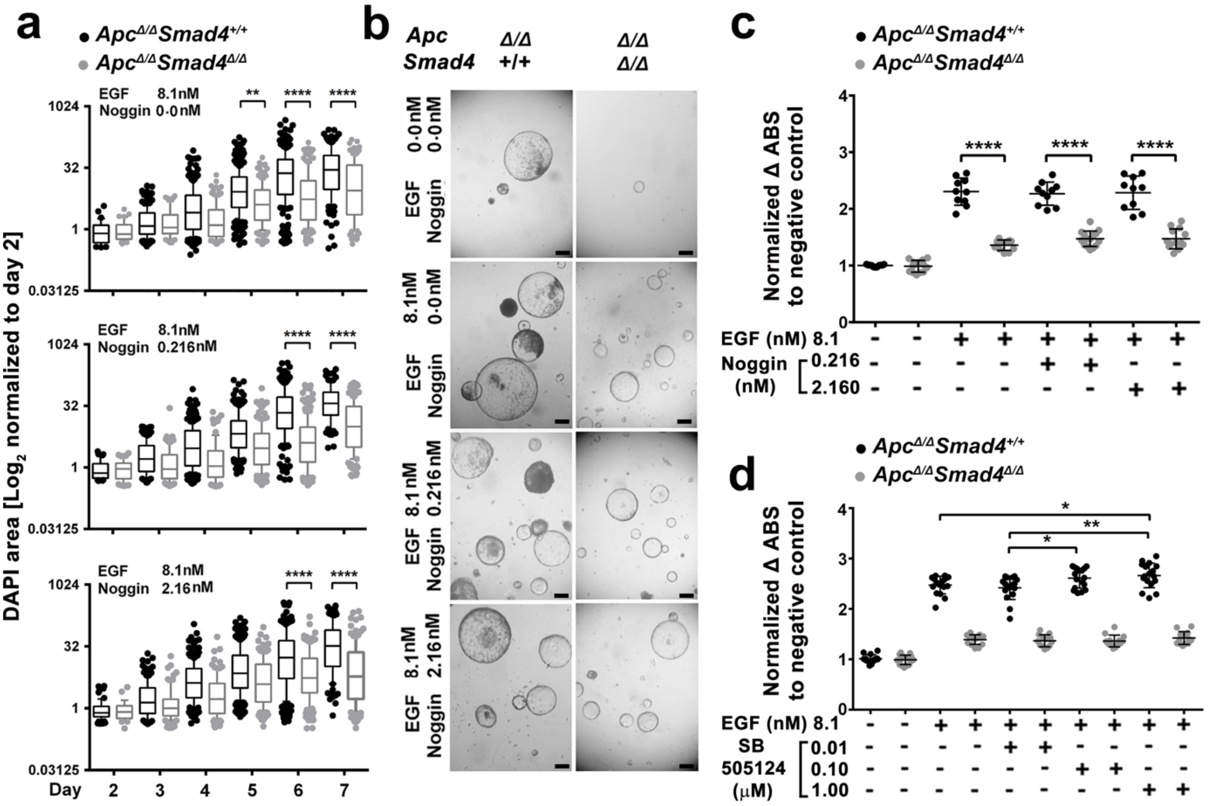


### Supplementary Figure 2

**Growth of *Apc*$^{\Delta/\Delta}$*Smad4*$^{\Delta/\Delta}$ adenoma are Noggin and SB-505124 independent**.

**a**. Adenomas were seeded at cell density of 5000 cells/well and adenoma growth measured either by cross-sectional area (DAPI) or by almar-Blue (AB), and tested against a 10-fold range in Noggin (0.216-2.16nM) and SB-505124 (0.01-1.0µM) in the presence of 8.1nM EGF (n=3 mice/group, 50 adenoma each). Y-axis $Log_2$, whiskers represent 10-90 percentile. One-way ANOVA followed by Sidak's multiple comparison test. **, **** indicates P<0.01 and P<0.0001, respectively. **b.** Bright field images showing adenomas from both genotypes on D7. Scale bars 100µm. **c, d.** Adenoma growth on D3 as measured by AB reduction and normalized to the negative control (Matrigel only). Adenomas were extracted by TryplE Express. (from n=2 *Apc*$^{fl/fl}$*Smad4*$^{+/+}$ mice and n=3 *Apc*$^{fl/fl}$*Smad4*$^{fl/fl}$ genotyped mice, 4-5 replicates in each experiment. One way-ANOVA with Tukey's multiple comparison test, *, **, ***, **** indicates P<0.05, P<0.01, P<0.001, P<0.0001, respectively.

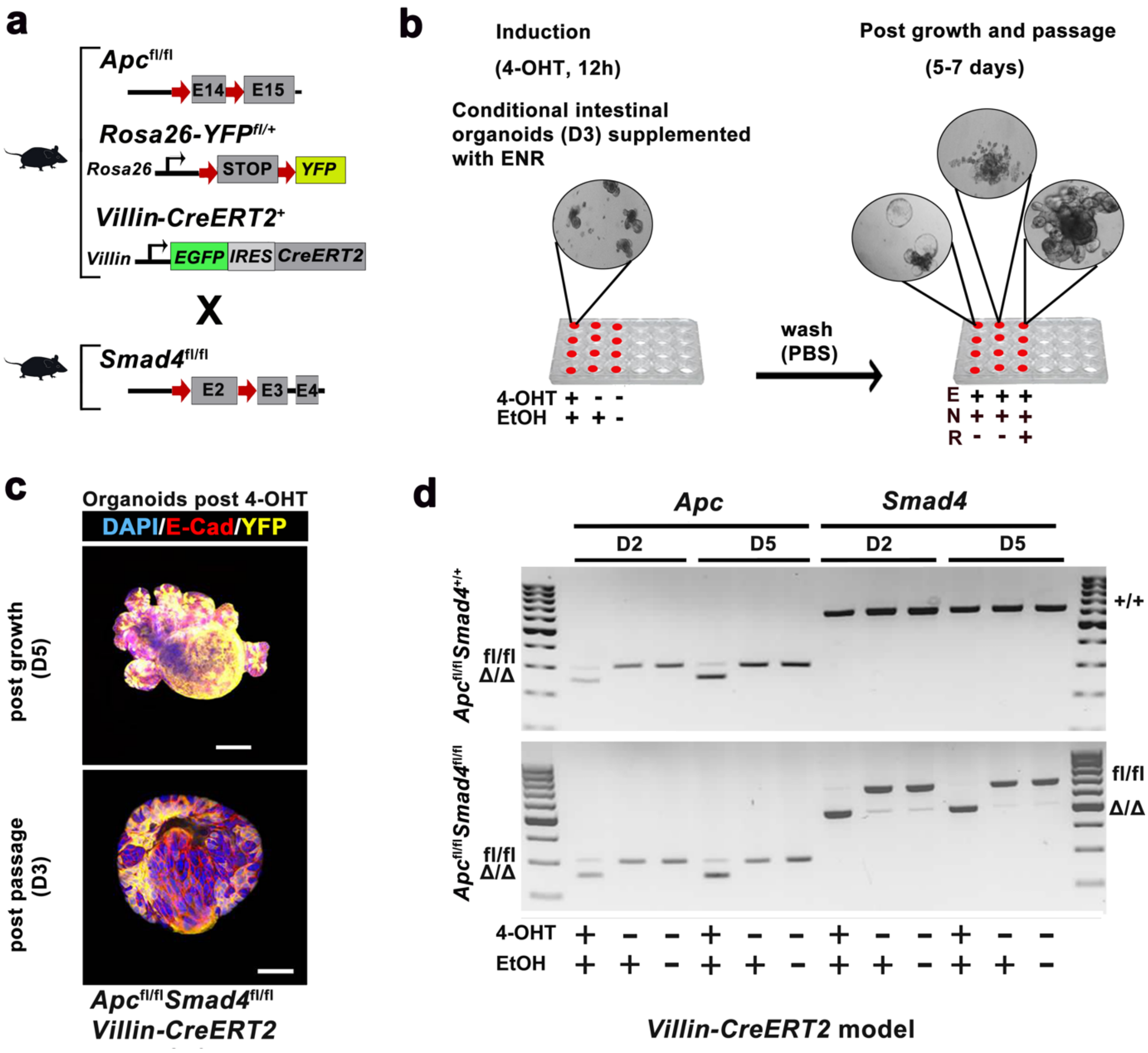


**Supplementary Figure 3**
**Conditional generation of crypt derived adenoma organoids using Villin Cre-ER$^{T2}$**

**a.** Simplified diagram representing the transgenic breeding strategy to generate experimental mice. **b.** Experimental growth requirement for conditional intestinal organoid manipulation with 4-OHT to generate intestinal adenomas *in vitro.* **c.** Confocal images of organoids showing the expression of Rosa26-YFP reporter post 4-OHT treatment in the *Apc*$^{Δ/Δ}$*Smad4*$^{Δ/Δ}$*Villin-CreERT2* manipulated organoids. Scale bars 50µm. **d.** PCR products show recombination of *Apc* (left side of both gels, lane 2, 5) and *Smad4* (right side lower gel, lane 8, 11) in *Apc, Smad4 Villin-CreERT2* adenoma model (genetically manipulated *in vitro* formed) on D2 and D5 post 4-OHT treatment (n=2 with similar results).

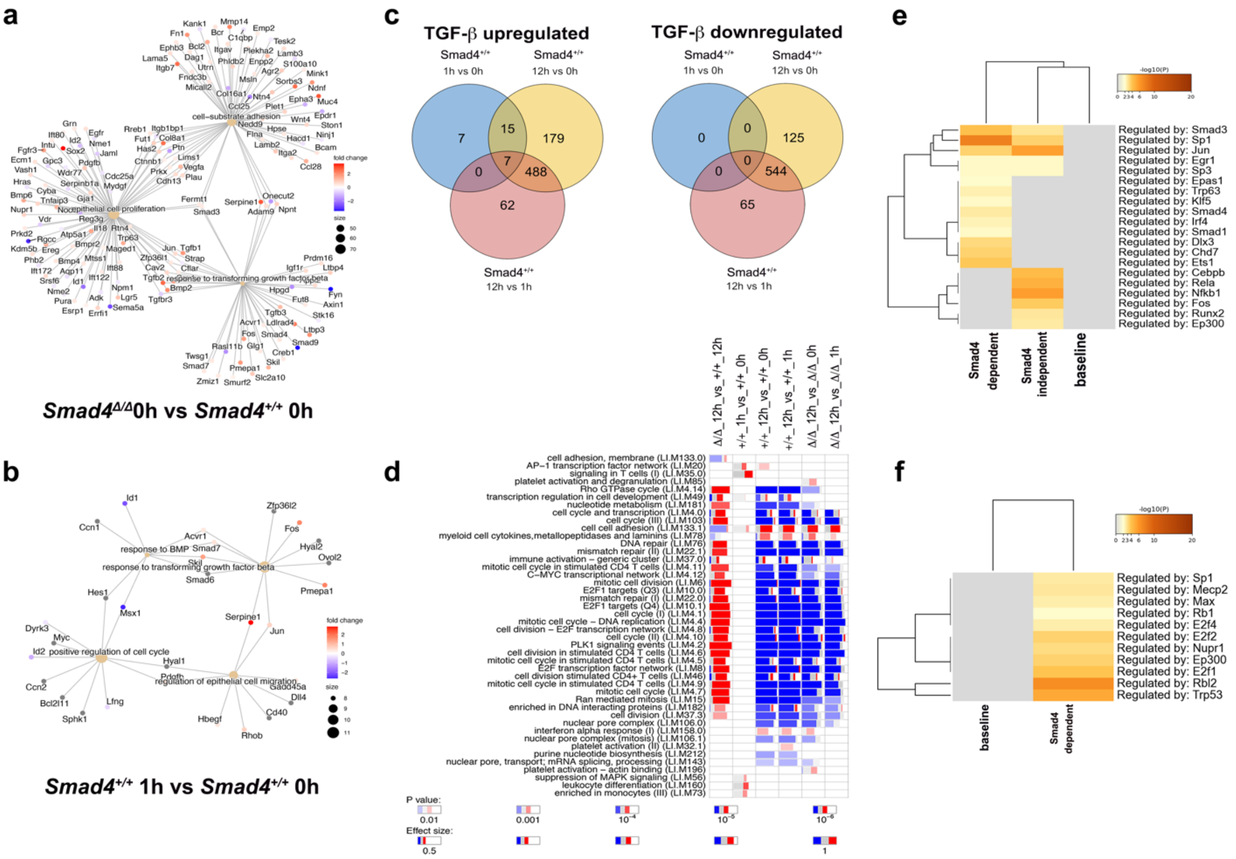


**Supplementary Figure 4**
**Enrichment analysis for TGF-β/Smad4 dependent genes.**

**a.** Pathway cnet plot for differentially regulated genes between *Apc*$^{\Delta/\Delta}$*Smad4*$^{+/+}$ and *Apc*$^{\Delta/\Delta}$*Smad4*$^{\Delta/\Delta}$ adenoma prior to TGF-β1 exposure (T=0). **b**. Pathway cnet plot for differential regulated genes as in a. but for alone comparing wild-type *Apc*$^{\Delta/\Delta}$*Smad4*$^{+/+}$ at t=0 and t=12 hr. **c**. Venn diagram showing upregulate and downregulated genes per wild-type *Apc*$^{\Delta/\Delta}$*Smad4*$^{+/+}$ genotype at different time points for (log2FC >1.5 and <-1.5, adjusted p-value), respectively. **d**. Enriched gene modules of DEG using the LI module gene set enrichment (GSEA with tmod package). See Fig. 3e for the Hallmark module analysis. Comparisons are shown in each column. Red and blue indicate the proportion of genes in a module that are either upregulated or downregulated, respectively. The width of each box relates to its effect size, while lighter, less saturated colours indicate lower significance with p-values. **e** and **f**. Heatmaps showing enriched transcription factors for Smad4 dependent and independent using TRRUST implemented in Metascape for e. upregulated and f. downregulated genes.

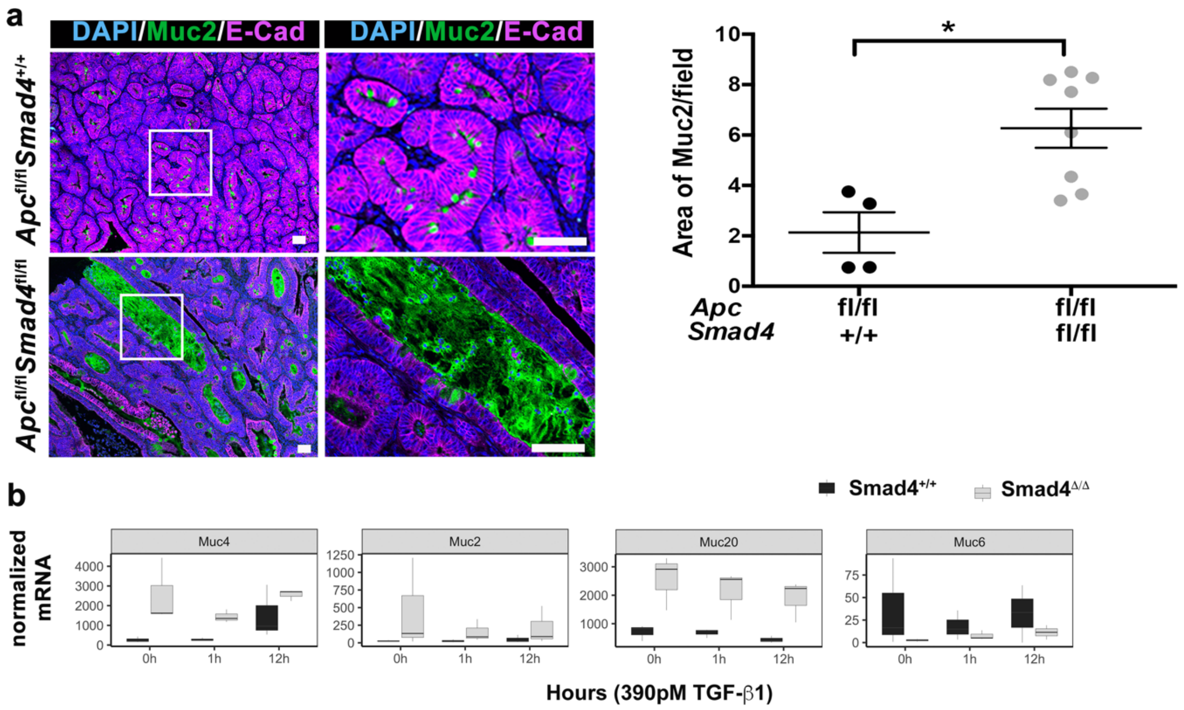


**Supplementary Figure 5**
**Smad4 dependent expression of Mucin genes**

**a.** Representative images of Muc2 labelling from caecal tissue sections of both *Apc*$^{fl/fl}$,*Smad4*$^{+/+}$ and *Apc*$^{fl/fl}$,*Smad4*$^{fl/fl}$ genotypes. White boxed regions are magnified below each one, scale bars 100µm. Quantification of Muc2 area in the sections by ImageJ represented by the percentage of Muc2 positive area to the tumour area. Statistical analysis was performed using student's t-test,*p=0.0153. Error bars±S.D. Each dot represents the average of area from 5 fields from sections of the same mouse (n=4 for *Smad4*$^{+/+}$, n=8 for *Smad4*$^{fl/fl}$). **b.** Expression of different mucin genes (*Muc4, Muc2, Muc20, Muc6*) from RNA-seq from normalised data.

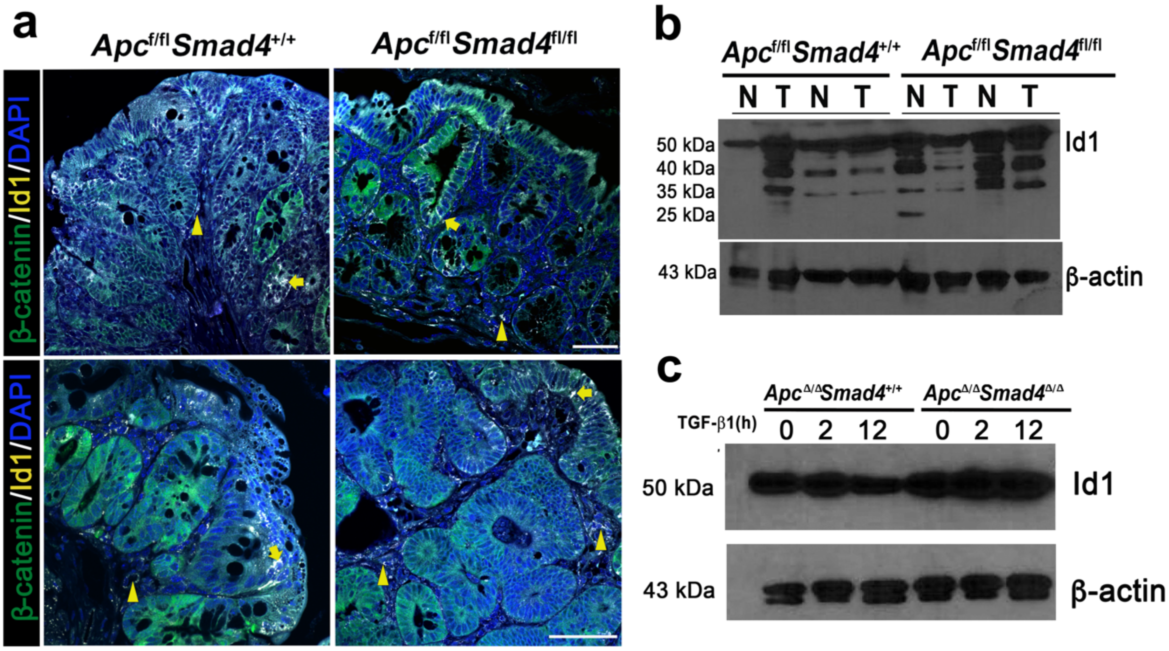


**Supplementary Figure 6**

***Id1* expression in *Apc$^{fl/fl}$Smad4$^{fl/fl}$Lgr5-Cre$^{ERT2}$* and *Apc$^{fl/fl}$Smad4$^{+/+}$Lgr5-Cre$^{ERT2}$* tissues and adenomas.**

**a.** Representative immunolabelling of adenoma tissue sections from *Apc*$^{fl/fl}$*Smad4*$^{+/+}$ and *Apc*$^{fl/fl}$*Smad4*$^{fl/fl}$ with an anti-Id1 antibody. The arrows show cytoplasmic Id1 in the epithelial tissues, and the arrowheads show cytoplasmic Id1 in the stroma of both genotypes. Scale bar 50µm. **b.** Western blot of Id1 (50kDa) from adenoma (T) and adjacent normal tissue (N) from *Apc*$^{fl/fl}$*Smad4*$^{+/+}$ and *Apc*$^{fl/fl}$*Smad4*$^{fl/fl}$. β-actin (43kDa) is a loading control. **c.** Western blot of Id1 from in *Smad4*$^{\Delta/\Delta}$ and *Smad4*$^{+/+}$ adenoma organoids at t=0, t=2 and t=12hr of 390pM TGF-β1 treatment.

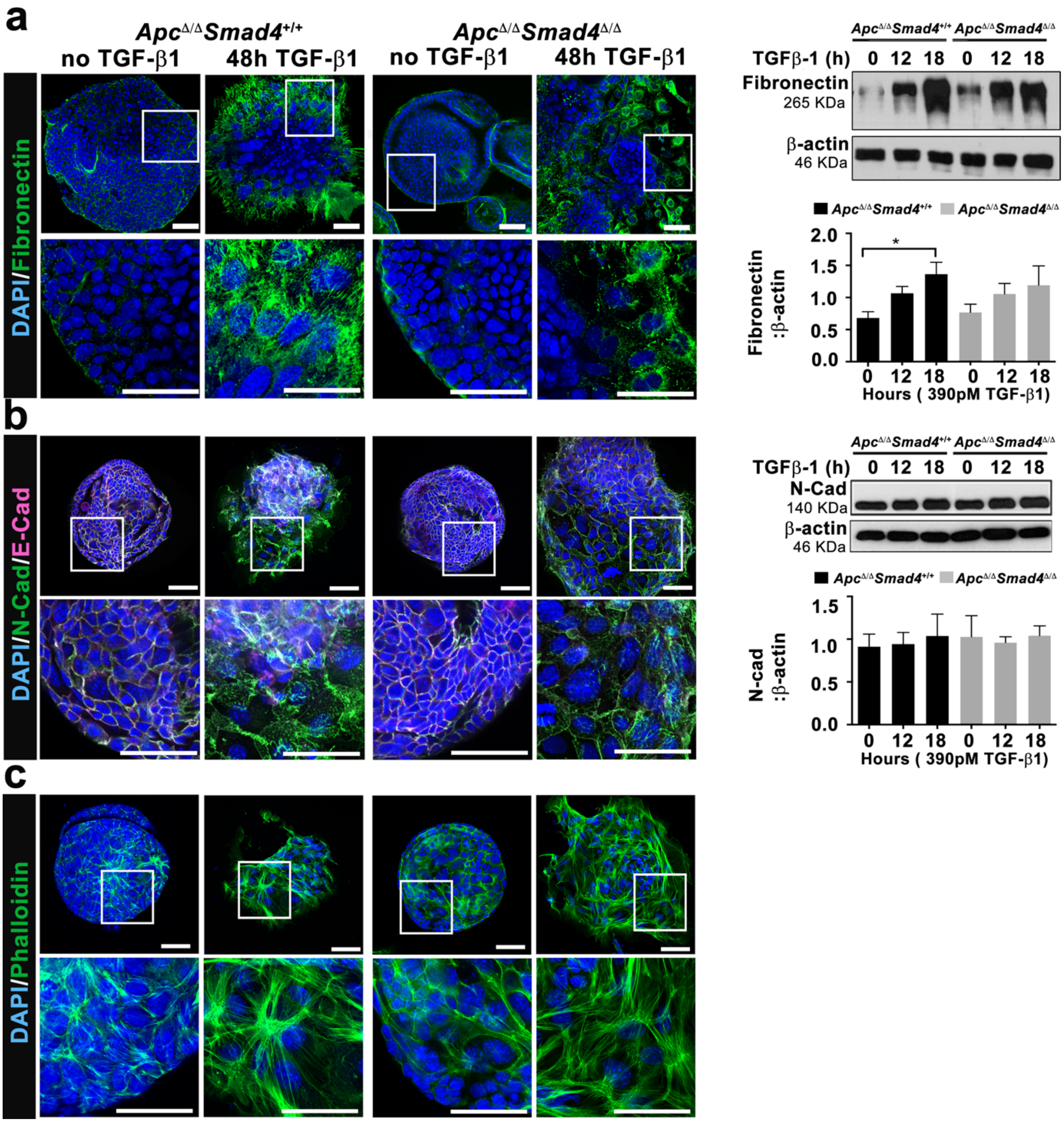


**Supplementary Figure 7**
**Smad4 dependent expression and localisation of epithelial-mesenchymal transition proteins**

**a.** Representative confocal images show *Smad4* independent upregulation of fibronectin (green) in adenomas post 390pM TGF-β1 treatment (left). Representative western blot for fibronectin and the ratio to β-actin as determined by densitometric analysis (n=3), *p=0.0429 (right). **b.** Downregulation of E-cadherin (magenta) and upregulation of N-cadherin (green) 48h post 390pM TGF-β1 treatment (left). Representative western blot for N-cadherin (and the ratio of it to β-actin as determined by densitometric analysis (n=3) (right). **c.** F-actin (by phalloidin) labelling 48h post 390pM TGF-β1 treatment. The untreated cells show a peripheral band of actin filaments, polarized epithelial cells and stress fibres in the treated adenomas. White boxed regions are magnified. Scale bars 50µm. Statistical analysis nonparametric one-way ANOVA with Dunn's multiple comparison test. Error bars±S.D. TGF-β1 concentration 390pM.

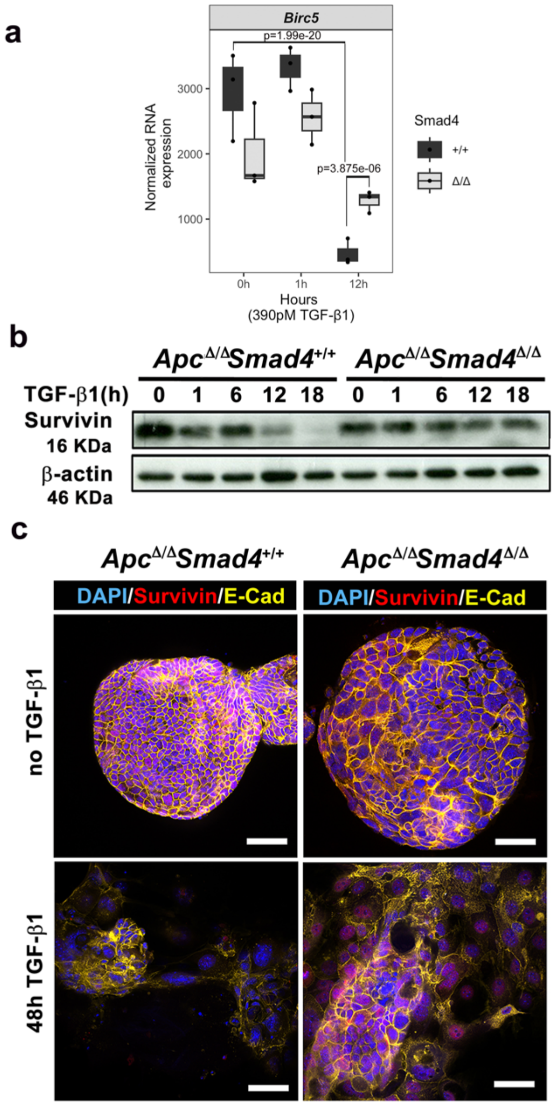


**Supplementary Figure 8**
**Smad4 dependent effects on *Birc5* (Survivin) expression with TGF-β1 treatment**.

**a.** Normalized *Birc5* (Survivin) expression from the RNA-seq taken at 0h, 1h and 12h following TGF-β1 exposure in Smad4$^{+/+}$ adenomas (black) and Smad4$^{\Delta/\Delta}$ (gray). Wald test followed by Benjamini-Hochberg for multiple hypothesis correction. **b.** Representative immunoblots showing the downregulation of Survivin within 12h TGF-β1 treatment predominantly in Smad4$^{+/+}$ adenomas. **c.** Confocal images showing Survivin nuclear localization after 48h of TGF-β1 treatment in Smad4$^{\Delta/\Delta}$ adenomas. Scale bars 50μm.

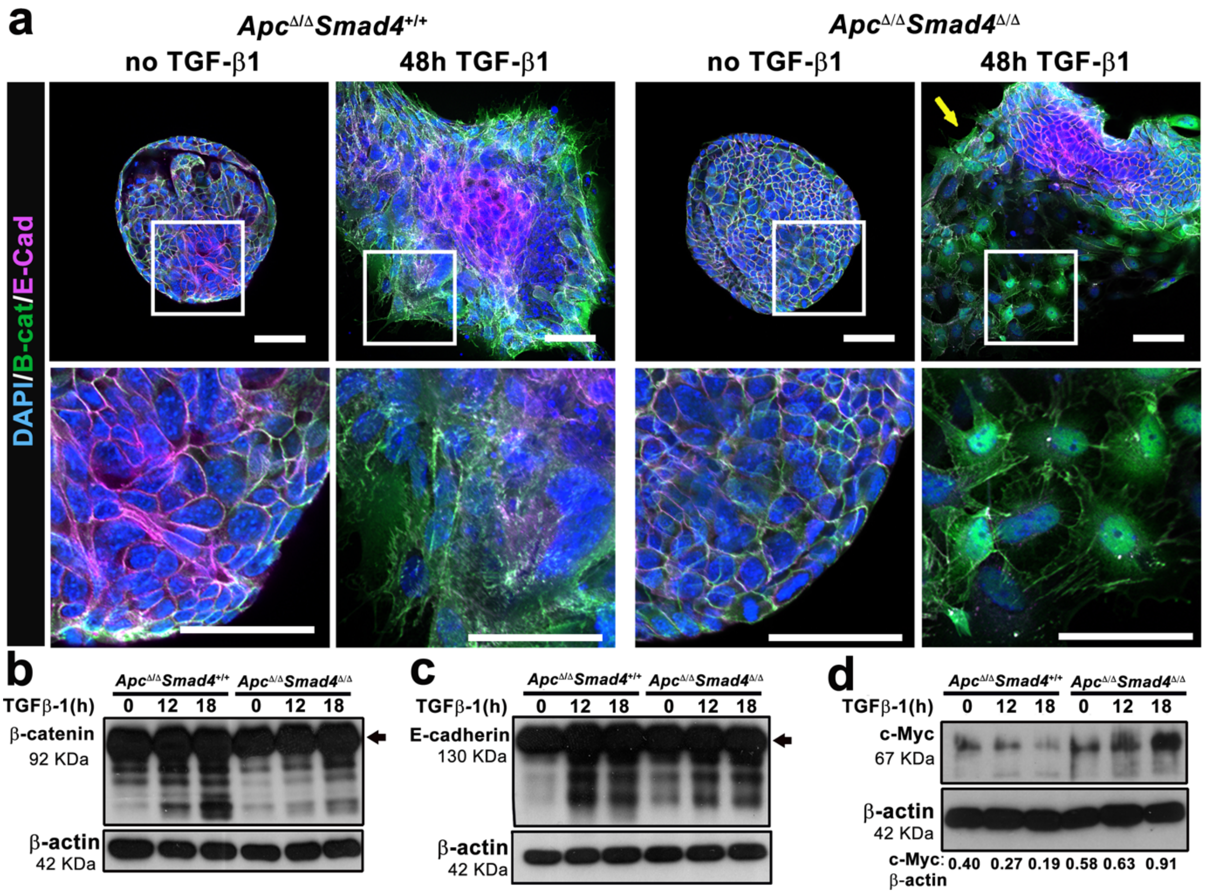


**Supplementary Figure 9**
**Smad4 dependent modification of the Wnt pathway**

**a.** Cytoplasmic and nuclear β-catenin and E-cadherin localisation following adenoma organoid TGF-β1 exposure. Representative confocal images show nuclear (DAPI) and cytoplasmic β-catenin with E-cadherin loss from the cytoplasmic cell membrane (yellow arrow) in the transformed adenoma post-TGF-β1. Magnification white-boxed regions are shown demonstrating nuclear β-catenin. Scale bars 50μm. **b, c, d.** Representative western blots showing **b.** β-catenin, **c.** E-cadherin and **d.** c-Myc at different time points following TGF-β1 exposure. β-catenin at 92 kDa (arrow) with multiple bands suggesting degradation. E-cadherin at 130 KDa (arrow), with multiple bands suggesting degradation. c-Myc at 67 KDa. The ratio of c-Myc to β-actin is represented below as determined by densitometric analysis (n=2). TGF-β1 concentration 390pM.

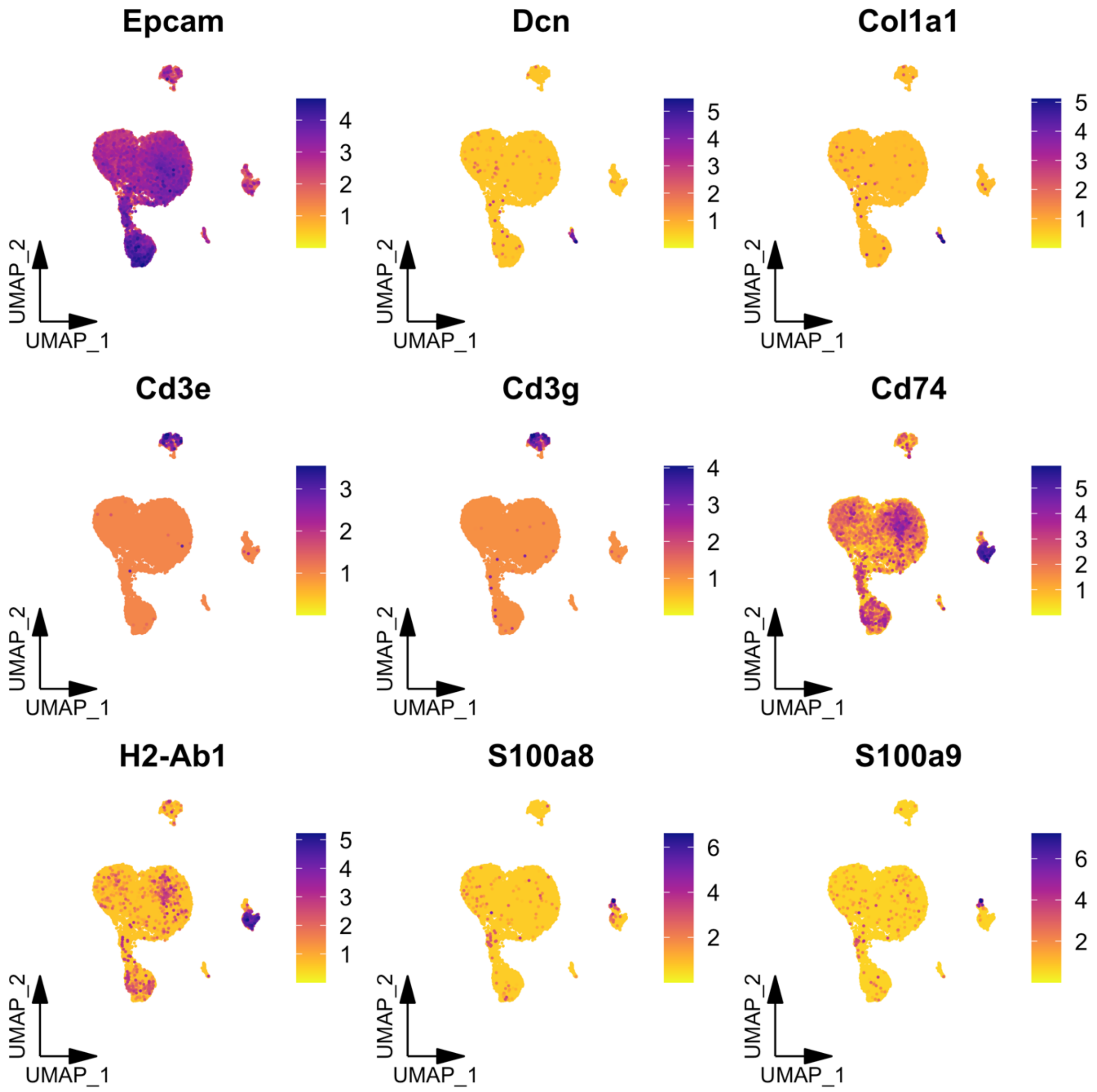


**Supplementary Figure 10**
**Single cell RNA-seq clusters of cell types in murine caecal tumours**

Unsupervised graph clustering using Seurat separated the cells into 20 clusters at a granularity of 1.2. These were visualised using UMAP and labelled according to the expression of known marker genes. The UMPAs show the projection of marker genes for epithelial cells (Epcam), cancer associated fibroblast (CAF) (DCN, Col1a1), T cells (Cd3e, Cd3g), Macrophages (Cd74, H2-Ab1, lyz2), and Neutrophils (S100a8 and S100a9).

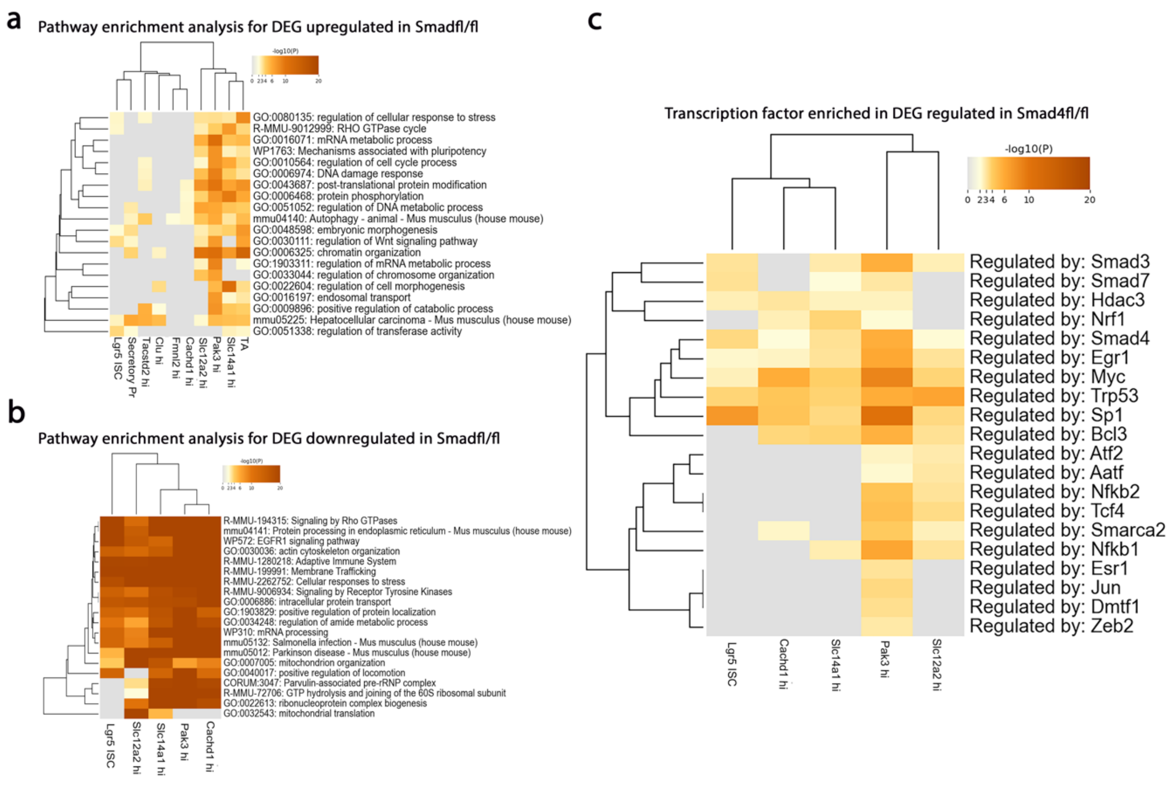


**Supplementary Figure 11**
***Smad4* dependent pathway enrichment analysis of single cell RNA-seq for different clusters**

**a.** and **b.** Heat maps of differentially expressed genes using GO ontology in *Apc*$^{fl/fl}$*Smad4*$^{fl/fl}$ that are **a**. upregulated and **b**. downregulated, data generated using Metascape. **c**. Heatmap showing enriched transcription factors for *Smad4* dependent upregulated genes using TRRUST implemented in metascape. Grey regions indicate non-significant differences.

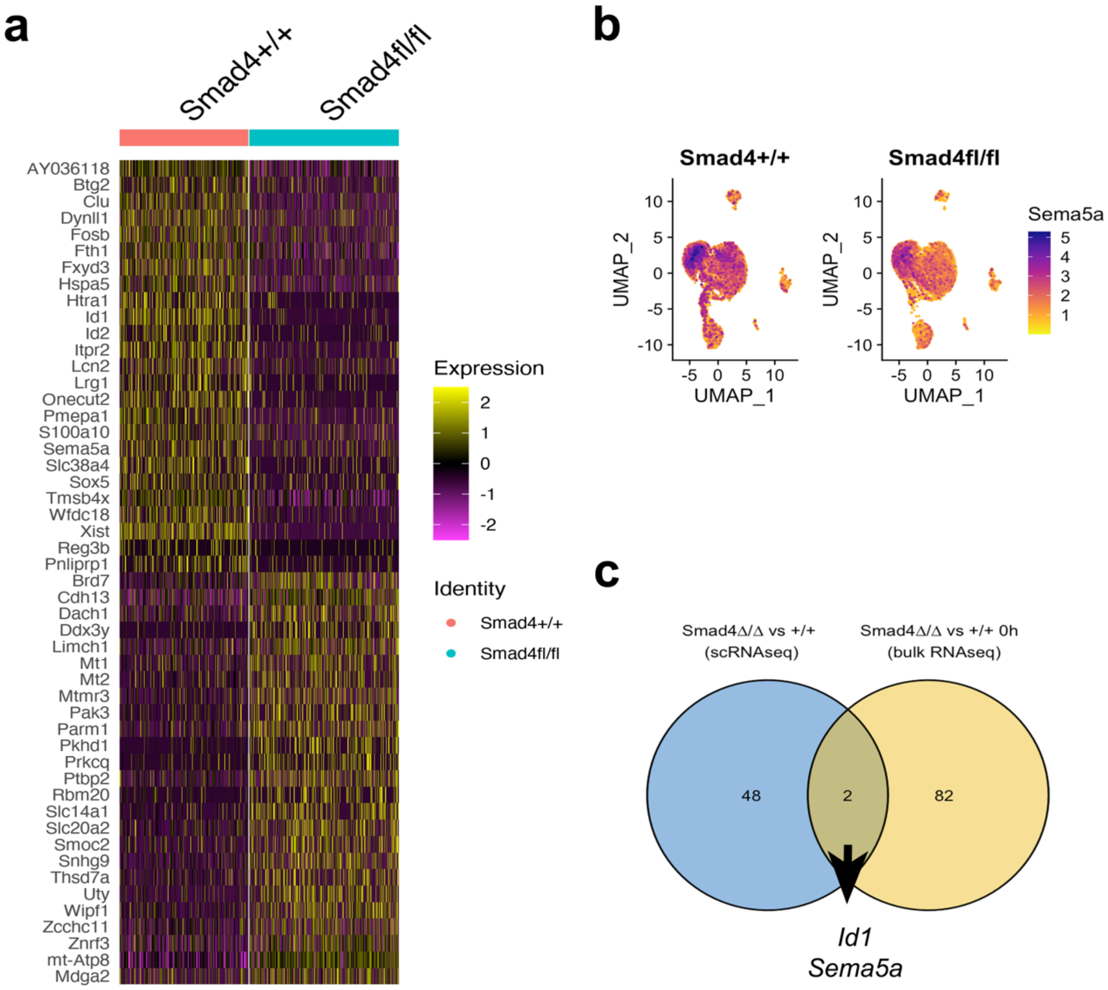


**Supplementary Figure 12**
***Smad4* dependent supplementary analysis of single cell RNA-seq**

**a**. Heat map of differentially expressed genes (top and bottom 50 genes) without clustering (pseudo-bulk analysis) between *Apc*$^{fl/fl}$*Smad4*$^{+/+}$ and *Apc*$^{fl/fl}$*Smad4*$^{fl/fl}$ derived caecal adenoma. Xist expression reflects unequal sex distribution between genotypes. Note the suppression of *Id1*, *Id2*, *Sema5a* and increased expression of *Pak3* and *Slc14a1*. **b.** UMAP feature plots for *Sema5a* between genotypes. **c**. Venn diagram of differential gene expression (log2FC >2 or <2) for bulk RNAseq from organoids (adjusted p value <0.05) resulted in 84 DEGs. This was compared to scRNA-seq DEGs for the top 25 and bottom 25 ranked gene. The only genes that overlapped between the two DEG groups were *Id1* and *Sema5a*.

**Supplementary Movies**

**Movie 1**: Wild-type adenoma organoids untreated
**Movie 2**: Wild-type adenoma organoids treated with TGF-β1
**Movie 3**: Smad4 null adenoma organoids untreated
**Movie 4**: Smad4 null adenoma organoids treated with TGF-β1

Movies from representative adenoma organoids of respective genotypes were cultured and images captured as described in Methods. Organoids were stabilised prior to imaging and TGF-β1 addition (390pM).
