## Supplementary Material and Methods for "*Smad4* and TGF-β1 dependent gene expression biomarkers in conditional intestinal adenoma, organoids and colorectal cancer"

| **KEY RESOURCES TABLE** |  | | | | |  | | | |
| --- | --- | --- | --- | --- | --- | --- | --- | --- | --- |
| **REAGENT or RESOURCE** | | **SOURCE** | | | **IDENTIFIER** | | | | |
| **Antibodies** | |  | | |  | | | | |
| Mouse IgG1 anti-β-catenin | | | | BD Bioscience | | | | | 610154, RRID:AB_397555 |
| Rabbit anit-CD11b | | | | Abcam | | | | | ab75476, RRID:AB_1310048 |
| Rabbit anti-CD3 | | | | Abcam | | | | | ab5690, RRID:AB_305055 |
| Rabbit anti-C-myc | | | | Abcam | | | | | ab11917, RRID:AB_298699 |
| Rabbit anti-Cleaved caspase3 (Asp175) (5A1E) | | | | Cell Signaling | | | | | 9664, RRID:AB_2070042 |
| Mouse IgG2a anti- E-cadherin | | | | BD Bioscience | | | | | 610182, RRID:AB_397581 |
| Rabbit anti-Fibronectin | | | | Abcam | | | | | ab23750, RRID:AB_447655 |
| Rat IgG2b anti-F4/80 (Cl:A3-1) | | | | Serotech | | | | | MCA497RT, RRID:AB_1102558 |
| Chicken IgY anti-GFP | | | | Abcam | | | | | ab13970, RRID:AB_300798 |
| Rabbit anti-Ki67 (SP6) | | | | Thermo Scientific | | | | | RM-9106-S0, RRID:AB_2341197 |
| Rabbit IgG anti-Mucin 2 (H-300) | | | | Santa Cruz | | | | | sc-15334,RRID:AB_2146667 |
| Rabbit anti-N-cadherin (EPR1792Y) | | | | Millipore | | | | | 04-1126, RRID:AB_1977064 |
| Rabbit IgG anti-Smad2 (D43B4) XP | | | | Cell Signaling | | | | | 5339, RRID:AB_10626777 |
| Rabbit IgG anti-Smad3 (C67H9) | | | | Cell Signaling | | | | | 9523, RRID:AB_2193182 |
| Mouse IgG1Smad4 (B-8) | | | | Santa Cruz | | | | | sc-7966, RRID:AB_627905 |
| Rabbit anti-Survivin (71G4B7) | | | | Cell Signaling | | | | | 2808S, RRID:AB_2063948 |
| Mouse IgG2ak Id1 (B-8) | | | | Santa Cruz | | | | | sc-133104, RRID: AB_2122863 |
| Rabbit anti-p-Smad2 (Ser465/467) | | | | Millipore | | | | | AB3849, RRID:AB_177440 |
| Rabbit anti-p-Smad3 (Ser423/425) (D12E11) | | | | Cell Signaling | | | | | 8769 |
| Rabbit anti-𝛼-tubulin | | | | Cell Signaling | | | | | 2144, RRID:AB_2210548) |
| Rabbit β-actin | | | | Abcam | | | | | ab8227, RRID:AB_2305186 |
| AF-555 goat anti-mouse IgG2a | | | | Invitrogen | | | | | A21137, RRID:AB_2535776 |
| AF-647 goat anti-mouse IgG1 | | | Invitrogen | | | | | A21240, RRID:AB_2535809 | |
| AF-647 goat anti-mouse IgG2a | | | Invitrogen | | | | | A21241, RRID:AB_2535810 | |
| AF-647 goat anti-rabbit IgG | | | Invitrogen | | | | | A21245, RRID:AB_2535813 | |
| AF-594 goat anti-mouse IgG2a | | | Invitrogen | | | | | A21135, RRID:AB_2535774 | |
| Goat pAb to chk IgY | | | Abcam | | | | | ab96952, RRID:AB_308760 | |
| Biotinylated goat anti-rabbit | | | Dako | | | | | E0432, RRID:AB_2313609 | |
| Biotinylated goat anti-mouse | | | Dako | | | | | E0433, RRID:AB_2687905 | |
| Biotinylated rabbit anti-rat | | | Dako | | | | | E0468 | |
| goat anti-rabbit horseradish peroxidase (HRP) | | | Dako | | | | | P0448, RRID:AB_2617138 | |
| goat anti-mouse HRP | | | Dako | | | | | P0447, RRID:AB_2617137 | |
| **Chemicals, peptides, and recombinant proteins** | | | | | | | | | |
| Advanced DMEM/F12 media (ADF) | Invitrogen | | | | | 12634 | | | |
| Growth factor reduced Matrigel | BD Bioscience | | | | | 536231 | | | |
| N2 | Invitrogen | | | | | 17502 | | | |
| B27 | Invitrogen | | | | | 12587 | | | |
| Epidermal growth factor (EGF) | Peprotech | | | | | AF-100-15 | | | |
| Bovine serum albumin (BSA) | Sigma | | | | |  | | | |
| Y-27632, ROCK Inhibitor | Tocris | | | | | 1254 | | | |
| SB-505124, TGF-β inhibitor | Sigma | | | | | S4696 | | | |
| Noggin (N) | Peprotech | | | | | | 250-38 | | |
| mR-spondin | R&D Systems | | | | | | 3474-RS | | |
| Tamoxifen | Sigma | | | | | | T5648 | | |
| **Experimental models: Organisms/strains** |  | | | | | |  | | |
| *Apc*^fl/fl^ (*Apc*^tm1Tno^) | <https://pubmed.ncbi.nlm.nih.gov/9311916/> | | | | | MGI:3764940 | | | |
| *Smad4*^fl/fl^ (*Smad4*^tm1Rob^) | <https://pubmed.ncbi.nlm.nih.gov/15215210/> | | | | | MGI:3624262 | | | |
| *RosaYFP*^fl/fl^ (Gt(ROSA)^26Sortm1^(EYFP)Cos) | <https://bmcdevbiol.biomedcentral.com/articles/10.1186/1471-213X-1-4> | | | | | MGI: 2449038 | | | |
| *Lgr5*^tm1(cre/ERT2)Cle^ (129P2/OlaHsd) | <https://pubmed.ncbi.nlm.nih.gov/17934449>/ | | | | | MGI:3764660 | | | |
| Vil-CreER^T2^ (Tg(Vil-cre/ERT2)23Syr) | <https://pubmed.ncbi.nlm.nih.gov/15282745/> | | | | | MGI:3053826 | | | |
| Critical commercial assays | | | | | | | | | |
| AlamarBlue | AbD Serotec | | | | | BUF012B | | | |
| Quick-RNA MiniPrep | Zymo Research | | | | | R1054 | | | |
| RNA Clean & concentrator^TM^-5 | Zymo Research | | | | | R1015 | | | |
| **Deposited data** | | | | | | | | | |
| RNA-seq COAD-READ | The Cancer Genome Atlas Network, 2012 | | | | | http://firebrowse.org/?cohort=COADREAD | | | |
| Survival data for the COAD-READ TCGA | https://www.sciencedirect.com/science/article/pii/S0092867418302290?via%3Dihub#tbl1 | | | | |  | | | |
| RNA-seq | This study | | | | | To be added on acceptance | | | |
| scRNAseq | This study | | | | | To be added on acceptance | | | |
| **Software and algorithms** | | | | | | | | | |
| ImageJ 1.48 software | https://imagej.net/Fiji | | | | | RRID: SCR_003070 | | | |
| Photoshop (CS5) | https://www.adobe.com/products/photoshop.html | | | | | RRID:SCR_014199 | | | |
| Prism version 6.0. | [https://www.graphpad.com](https://www.graphpad.com/) | | | | | SCR_000306 | | | |
| DESeq2 version 1.28 | https://bioconductor.org/packages/release/bioc/html/DESeq2.html | | | | | RRID: SCR_015687 | | | |
| R version 4.0.2 & version 4.2.3 | <http://www.r-project.org/> | | | | | RRID:SCR_001905 | | | |
| TopHat aligner version 2.1.0 | https://ccb.jhu.edu/software/tophat/index.shtml | | | | | RRID:SCR_013035 | | | |
| Subread version 1.6.0 | http://subread.sourceforge.net | | | | | RRID:SCR_009803 | | | |
| clusterProfiler v3.18.1 | https://bioconductor.org/packages/release/bioc/html/clusterProfiler.html | | | | | RRID:SCR_016884 | | | |
| Survminer v0.4.9 | https://cran.r-project.org/web/packages/survminer/readme/README.html | | | | |  | | | |
| cBioPortal | https://www.cbioportal.org | | | | | RRID:SCR_014555 | | | |
| Cell Ranger Software Suite (v 7.0.1) | https://www.10xgenomics.com/support/software/cell-ranger/latest/release-notes/cr-release-notes | | | | | RRID:SCR_017344 | | | |
| scDblFinder (v1.12.0) | https://github.com/plger/scDblFinder | | | | | RRID:SCR_022700 | | | |
| Seurat (v4.3.0) | https://satijalab.org/seurat/ | | | | | RRID:SCR_016341 | | | |
| clustree (v0.5.1) | https://cran.r-project.org/web/packages/clustree/index.html | | | | | RRID:SCR_016293 | | | |
| scProportionTest | https://github.com/rpolicastro/scProportionTest | | | | | NA | | | |
| Metascape | https://scicrunch.org/resolver/RRID:SCR_016620?q=&i=rrid:scr_016620 | | | | | RRID:SCR_016620 | | | |
