## Supplementary Tables for "*Smad4* and TGF-β1 dependent gene expression biomarkers in conditional intestinal adenoma, organoids and colorectal cancer"

Excel Files

Supplementary Table 1

1. Supplementary Table 1a: DEG in Smad4∆/∆ 0h compared to Smad4+/+ 0h.
2. Supplementary Table 1b: DEG in Smad4∆/∆ 1h compared to Smad4+/+ 1h.
3. Supplementary Table 1c: DEG in Smad4∆/∆ 12h compared to Smad4+/+ 12h.
4. Supplementary Table 1d: DEG in Smad4+/+ 1h compared to Smad4+/+ 0h.
5. Supplementary Table 1e: DEG in Smad4+/+ 12h compared to Smad4+/+ 0h.
6. Supplementary Table 1f: DEG in Smad4+/+ 12h compared to Smad4+/+ 1h.
7. Supplementary Table 1g: DEG in Smad4∆/∆ 12h compared to Smad4∆/∆ 0h.
8. Supplementary Table 1h: DEG in Smad4∆/∆ 12h compared to Smad4∆/∆ 1h.
9. Supplementary Table 1i: Summary of Log_2_FC and adjusted p-value for the eight pairwise comparisons.

Supplementary Table 2.

1. Supplementary Table 2a: Pathway enrichment analysis using MSigDB Hallmark gene sets with tmod.
2. Supplementary Table 2b: Genes enriched in tmod analysis from MSigDB Hallmark gene sets.

Supplementary Table 3

1. Supplementary Table 3a: TGF-β Early response genes upregulated in Smad4+/+ adenomas (Log_2_FC and adjusted p-value).
2. Supplementary Table 3b: TGF-β late response genes upregulated in Smad4+/+ adenomas (Log_2_FC and adjusted p-value).
3. Supplementary Table 3c: TGF-β late response genes downregulated in Smad4+/+ adenomas (Log_2_FC and adjusted p-value).
4. Supplementary Table 3d: TGF-β Smad4 independent Genes (gene name, description, and direction).
5. Supplementary Table 3e: TGF-β Smad4 dependent Genes (gene name, description, and direction).
6. Supplementary Table 3f: Genes regulated at the baseline (gene name, description, and direction).

Supplementary Table 4

1. Marker gene for all clusters from the single cell RNA-seq at 1.2 resolution.
2. Marker gene for clusters 10 and 22 from the single cell RNA-seq at 2.2 resolution.

Supplementary Table 5

1. Supplementary Table 5a: Patients number in the study groups.
2. Supplementary Table 5b: Fold difference in the RSEM median expression of *ID1* between different groups.
3. Supplementary Table 5c: Fold difference in the RSEM median expression of *ID1*
4. between different groups with no BMP pathway mutation.
5. Supplementary Table 5d: Kruskal-Wallis p-values for genes evaluated in the TCGA cohorts in the context of Smad4 mutation.
6. Supplementary Table 5e: Multiple comparisons for genes with significant Kruskal-Wallis comparison (Dunn test).
7. Supplementary Table 5f: Significant genes in adenoma at 0h with absolute Log_2_FC >2 and adjusted p-value <0.05 and their human orthologs.
8. Supplementary Table 5g: Comparison of *ID1, SPP1, PAK3* expression between different groups in TCGA cohorts.
9. Supplementary Table 5h: TCGA cohorts with the mutation status used in the study.
